# FLAIRR-seq: A novel method for single molecule resolution of near full-length immunoglobulin heavy chain repertoires

**DOI:** 10.1101/2022.09.24.509352

**Authors:** Easton E. Ford, David Tieri, Oscar Rodriguez, Nancy Francoeur, Juan Soto, Justin Kos, Ayelet Peres, William Gibson, Catherine A. Silver, Gintaras Deikus, Elizabeth Hudson, Cassandra R. Woolley, Noam Beckmann, Alexander Charney, Thomas C. Mitchell, Gur Yaari, Robert P. Sebra, Corey T. Watson, Melissa L. Smith

## Abstract

Current Adaptive Immune Receptor Repertoire Sequencing (AIRR-seq) strategies resolve expressed antibody (Ab) transcripts with limited resolution of the constant region. Here we present a novel near full-length AIRR-seq (FLAIRR-Seq) method that utilizes targeted amplification by 5’ rapid amplification of cDNA ends (RACE), combined with single molecule, real-time sequencing to generate highly accurate (>Q40, 99.99%) IG heavy chain transcripts. FLAIRR-seq was benchmarked by comparing IG heavy chain variable (IGHV), diversity (IGHD), and joining (IGHJ) gene usage, complementarity-determining region 3 (CDR3) length, and somatic hypermutation to matched datasets generated with standard 5’ RACE AIRR-seq and full-length isoform sequencing. Together these data demonstrate robust, unbiased FLAIRR-seq performance using RNA samples derived from peripheral blood mononuclear cells, purified B cells, and whole blood, which recapitulated results generated by commonly used methods, while additionally resolving novel IG heavy chain constant (IGHC) gene features. FLAIRR-seq data provides, for the first time, simultaneous, single-molecule characterization of IGHV, IGHD, IGHJ, and IGHC region genes and alleles, allele-resolved subisotype definition, and high-resolution identification of class-switch recombination within a clonal lineage. In conjunction with genomic sequencing and genotyping of IGHC genes, FLAIRR-seq of the IgM and IgG repertoires from 10 individuals resulted in the identification of 32 unique IGHC alleles, 28 (87%) of which were previously uncharacterized. Together, these data demonstrate the capabilities of FLAIRR-seq to characterize IGHV, IGHD, IGHJ, and IGHC gene diversity for the most comprehensive view of bulk expressed Ab repertoires to date.

## Introduction

Antibodies (Abs) or immunoglobulins (IGs) are the primary effectors of humoral immunity and are found as both membrane-bound receptors on B cells and circulating, secreted proteins (1). Both membrane-bound B cell receptors (BCRs) and secreted Abs act to recognize and bind antigen. All Abs and BCRs are composed of two identical heavy and light chains that are post-translationally associated. The heavy chain is comprised of two distinct domains: (i) the variable domain (Fab), which allows for antigen binding, and (ii) the constant domain (Fc), which modulates downstream effector functions (1, 2). The light chain also includes a variable domain that, once post-translationally associated with the heavy chain variable domain, interacts with cognate antigen (3). In humans, Abs are grouped into discrete isotypes and subisotypes (i.e., IgM, IgD, IgG1, IgG2, IgG3, IgG4, IgA1, IgA2, and IgE), based on the expression of specific constant (C) genes within the IG heavy chain locus (IGH). Each isotype and subisotype has unique effector properties that together represent the wide diversity of Ab-mediated functions, including binding of Fc receptors (FCR), activation of complement, opsonization, antibody-dependent cellular cytotoxicity (ADCC) and antibody-dependent cellular phagocytosis (ADCP) (4, 5).

To facilitate the development of diverse Ab repertoires capable of recognizing the wide range of pathogens humans encounter, the IG genomic loci are highly polymorphic and harbor diverse and complex sets of genes that recombine in each B cell to encode up to 10^13^ unique specificities (6). B cells create this expansive catalog of specificities through somatic recombination of the variable (V), diversity (D), and joining (J) genes in IGH, and V and J genes from the corresponding light chain loci, lambda (IGL) and kappa (IGK) (7). During VDJ recombination in IGH, a single D and J gene are first recombined, while the intervening and unselected D and J gene sequences are removed by RAG recombinase (8). After D and J genes are joined, further recombination of a specific V gene to the DJ gene cassette completes the formation of the full VDJ rearrangement. Following transcription of the recombined VDJ, a single constant region gene is spliced together with the VDJ cassette to generate the completed heavy chain transcript (7). Recombination at IGL and IGK occurs similarly, recombining V and J genes only. Heavy and light chain transcripts are independently translated and linked via covalent cysteine bonds resulting in a fully functional protein prior to B cell cell-surface expression or secretion (Figure 1A) (9). Naïve B cells, which develop in the bone marrow from hematopoietic stem cell progenitors, have undergone VDJ recombination but have not yet encountered antigen, and solely express IgM and IgD (10). These naïve B cells then migrate to B cell zones in secondary lymphoid tissues where they encounter antigen, driving further maturation and class switch recombination (CSR) to enable the most effective humoral responses (11). CSR mediates the excision of IGHC genes at the DNA level, which leads to the utilization and linkage of different IGHC genes to the same VDJ, ultimately resulting in class switching to alternate isotypes and subisotypes (11).

**Figure 1.**
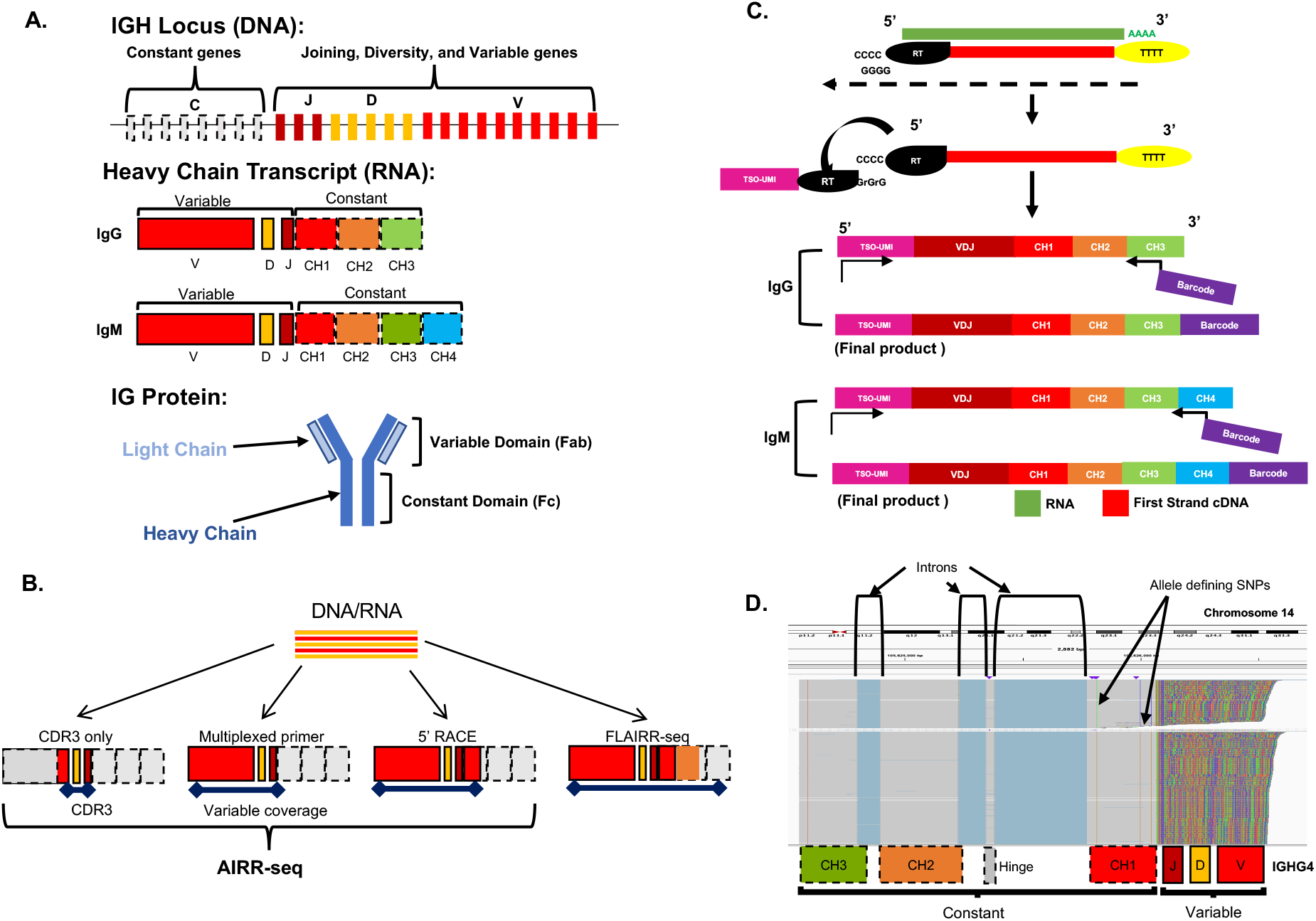
Overview of Ab structure and FLAIRR-seq molecular method. (A) Schematic representation of the IGH locus, heavy chain transcript structure, and functional IG protein. (B) Comparative coverage across the heavy chain transcript of commonly used AIRR-seq methods compared with FLAIRR-seq. (C) FLAIRR-seq molecular pipeline: RNA (brown) was converted to first-strand cDNA (red) using the 5’ RACE method, incorporating a 5’ TSO-UMI (pink) via template switch. Second strand amplification specifically targeted IgG and IgM molecules through priming of the 5’ TSO-UMI and the 3’ constant region IGH exon 3 (CH3) for IgG, or CH4 for IgM. A 16bp barcode was incorporated into the 3’ CH3/CH4 primers to enable sample multiplexing post-amplification. (D) IGV screenshot showing near full-length single molecule structure of IGHG4 FLAIRR-seq transcripts.

The IgG isotype class is represented by four subisotypes: IgG1, IgG2, IgG3, and IgG4. Each IgG subisotype circulates at varied frequencies and facilitates unique immune functions. For example, IgG1 is typically the most abundant circulating IgG and mediates proinflammatory responses; IgG2 targets bacterial polysaccharides, providing protection from bacterial pathogens; IgG3 confers protection against intracellular bacterial infections and enables clearing of parasites; and IgG4 contains exclusive structural and functional characteristics often resulting in anti-inflammatory and tolerance-inducing effects (5). Multiple studies have identified Ab-mediated subisotype-specific pathogenicity in the context of autoimmune diseases and cancer highlighting the need for further investigation of subisotype-specific repertoires (12-15).

Current Adaptive Immune Receptor Repertoire sequencing (AIRR-seq) methods aim to resolve variable and constant region transcripts to differing extents. Profiling of the variable region, even in part, defines V, D and J gene usage while also providing characterization of complementarity determining regions (CDR) 1, 2, and 3, which are hypervariable and directly interact with target antigen (Figure 1B) (16). CDR3-targeted profiling approaches, such as those used by Adaptive Biotechnologies (17), allow for V, D, and J gene assignments but do not provide complete resolution of the entire variable region (17). Multiplexed primer-based AIRR-seq strategies generate full variable region content but require specific primers to known targets and therefore may miss novel genes and alleles. 5’ RACE AIRR-seq methods capture the full-length VDJ exon without variable region-targeted multiplexed primer pools therefore limiting the impact of primer bias and enabling discovery of novel IGHV, IGHD, and IGHJ genes and alleles (18). 5’ RACE methods also often prime from the first IGHC exon (CH1), allowing for determination of isotype. Additional methods have been developed that shift amplification strategies by capturing additional IGHC region sequence to enable subisotype resolution; however, these methods sacrifice full and contiguous characterization of the IGHV gene (19). All commonly used AIRR-seq methods are further technically limited by the length restrictions (≤ 600nt) of short-read next generation sequencing. As a result, no current AIRR-seq strategy resolves the complete heavy chain transcript, including all IGHC exons alongside the recombined IGHV, IGHD, and IGHJ genes. Our team has recently shown that population-based polymorphisms within the IGHV, IGHD, IGHJ and IGL loci are far more extensive than previously known; the IGHC region has also been shown to contain genomic diversity, although the extent of this diversity has likely not been fully explored (20-27). Although it is understood that the Fc domain mediates Ab effector functions, there is limited knowledge as to how genetic variation in this region may impact functional capabilities or posttranslational modification (5, 28, 29). As such, there is a growing need to understand genomic variation across the complete Ab molecule. To address these limitations, we have developed novel, end-to-end pipeline to target, profile, and characterize the Ab heavy chain repertoire in the context of isotype (IgG, IgM) and subisotype (IgG1, IgG2, IgG3, IgG4).

Here, we present FLAIRR-seq, a novel targeted 5’ RACE-based amplification of near full-length IgG and IgM heavy chain transcripts, paired with single molecule real time (SMRT) sequencing, resulting in highly accurate (mean read accuracy ∼Q60, 99.9999%), near full-length Ab sequences from RNA derived from whole blood, isolated PBMC, and purified B cells. When analyzed with the Immcantation AIRR-seq tool suite (30, 31), we demonstrate that FLAIRR-seq performs comparably to standard 5’RACE AIRR-seq methods and single-molecule isoform sequencing (Iso-Seq) strategies for characterizing the expressed Ab repertoire. We further highlight the novel features of FLAIRR-seq data, including phased identification of IGHV, IGHD, IGHJ, and IGHC genes, facilitating the profiling of subisotype- and IGHC allele-specific repertoires and CSR characterization.

## Materials and Methods

### Sample collections

Experiments were conducted using healthy donor peripheral blood mononuclear cells (PBMC), purified B cells from healthy donors, or whole blood collected from hospitalized COVID-19 patients (Supplementary Table 1). Commercially available healthy donor PBMC (STEMCELL Technologies) and a subset of matched purified B cells were utilized to generate AIRR-seq and FLAIRR-seq validation datasets. Full-length isoform sequencing (Iso-Seq) was performed using B cells isolated from the PBMC of a healthy, consented 57-year-old male donor at the University of Louisville (UofL) School of Medicine. The UofL Institutional Review Board approved sample collection (IRB 14.0661). For COVID-19 affected patient samples (n=5), whole blood was collected from the Mount Sinai COVID-19 biobank cohort of hospitalized COVID-19 patients, approved by the Institutional Review Board at the Icahn School of Medicine at Mount Sinai as previously described (32).

### PBMC isolation and B cell purification

Frozen healthy donor PBMCs were purchased, thawed, and aliquoted for use in downstream experiments (STEMCELL Technologies). For Iso-Seq analyses, 175mL of venous blood was collected in a final concentration of 6mM K_3_EDTA using standard phlebotomy. PBMCs were isolated using Sepmate PBMC Isolation Tubes (STEMCELL Technologies) as previously described (33), with an additional granulocyte depletion step using the RosetteSep Human Granulocyte Depletion Cocktail (STEMCELL Technologies) as directed by the manufacturer. B cells from the freshly collected and frozen healthy donor PBMC were isolated using the EasySep Human Pan-B Cell Enrichment Kit, as described by the manufacturer (STEMCELL Technologies). Briefly, B cells, including plasma cells, were isolated by negative selection using coated magnetic particles. First, the B cell enrichment cocktail was added to the sample and mixed for a 5-minute incubation at room temperature, followed by addition of magnetic particles and further incubation for 5 minutes on the benchtop. The sample tube was then placed on an EasySep magnet (STEMCELL Technologies), and purified B cells were carefully eluted from the magnetic particles and immediately used for RNA extraction.

### Genomic DNA and RNA extraction

For the healthy frozen PBMC and matched purified B cells, genomic DNA (gDNA) and RNA were co-extracted using the AllPrep DNA/RNA Mini Kit (Qiagen) according to the manufacturer’s instructions. For the freshly processed UofL healthy donor PBMC, purified Pan-B cells were lysed in Buffer RLT Plus, and RNA was extracted with the RNeasy Plus Mini Kit (Qiagen) per the manufacturer’s protocol; no gDNA was collected from this sample. COVID-19 whole blood-derived RNA was extracted from samples collected in Tempus Blood RNA tubes using the Tempus Spin RNA Isolation Kit (ThermoFisher) as described by the manufacturer. For all samples, concentrations of RNA and gDNA (when appropriate) were assessed using the Qubit 4.0 fluorometer, with the RNA HS Assay Kit and Qubit DNA HS Assay Kit, respectively (ThermoFisher Scientific). RNA and gDNA integrity were evaluated using the Bioanalyzer RNA Nano Kit and DNA 1200 Kit, respectively (Agilent Technologies). Extracted RNA and gDNA were stored at -80°C and -20°C, respectively, until used.

### FLAIRR-seq targeted amplification of heavy chain transcripts

Extracted RNA was thawed on ice and converted to first strand complementary DNA (cDNA) using the SMARTer RACE 5’/3’ Kit (Takara Bio USA), as described by the manufacturer and a custom oligonucleotide that contained the template switch oligo and a unique molecular identifier (5’ TSO-UMI) for template switch during first strand cDNA synthesis. The following reaction conditions were used: (i) a primary master mix was prepared with 4.0 μL 5X First-Strand Buffer, 0.5 μL DTT (100 mM), and 1.0 μL dNTPs (20 mM) per reaction and set aside until needed; (ii) in a separate 0.2-mL PCR tube, 10 μL of sample RNA and 1 μL 5’-CDS Primer A were combined and incubated in a thermal cycler at 72°C (lid temperature: 105°C) for 3 minutes, followed by cooling to 42°C for 2 minutes; (iii) after cooling, tubes were spun briefly to collect contents and 1μL (12μM) of the 5’ TSO-UMI was added to the RNA; (iv) 0.5 μL of RNase inhibitor and 2.0 μL of SMARTScribe Reverse Transcriptase were added to the primary master mix tube per sample and 8 μL of the combined master mix was then added to each RNA-containing sample tube. First-strand cDNA synthesis reactions were incubated in a thermal cycler at 42°C (lid temperature: 105°C) for 90 mins, followed by heat inactivation at 70°C for 10 minutes. Total first strand cDNA generated in this reaction was diluted 1:2 with Tricine-EDTA Buffer before moving onto targeted heavy chain transcript amplification.

To specifically amplify heavy chain transcripts from total first-strand cDNA, targeted IgG and IgM transcript amplification reactions were performed using barcoded IgG (3’ primer binding in the constant region exon 3, CH3) or IgM (3’ primer binding in the constant region exon 4, CH4)-specific primers (Supplementary Table 2) and the following conditions: (i) 5 μL of diluted firststrand cDNA was added to 0.2-mL PCR tubes; (ii) a master mix was generated using 10 μL 5X PrimeSTAR GXL Buffer, 4 μL GXL dNTP mixture, 28 μL PCR-grade water, 1 μL PrimeSTAR GXL Polymerase and 1 μL 10x UPM form the SMARTer RACE 5’/3’ Kit per reaction; (iii) 44 μL master mix was added to each reaction tube followed by 1 μL of the appropriate barcoded IgG (CH3) or IgM (CH4) primer. Different temperatures were used for annealing of IgG- (63.5°C) and IgM-specific primers (60°C) to account for primer specific melting temperatures and to enhance targeted amplification specificity. Amplification conditions for full-length IgG were: 1 minute at 95°C, followed by 35 amplification cycles of 95°C for 30 sec, 63.5°C for 20 sec, and 2 minutes at 68°C, followed by a final extension for 3 minutes at 68°C and hold at 4°C. Amplification conditions for full-length IgM were: 1 minute at 95°C, followed by 35 amplification cycles of 95°C for 30 sec, 60°C for 20 sec., and 2 minutes at 68°C, followed by a final extension for 3 minutes at 68°C and hold at 4°C. Final amplification reactions were purified using a 1.1x (vol:vol) cleanup with ProNex magnetic beads (Promega). Successfully amplified products were quantified with Qubit dsDNA HS assay (ThermoFisher Scientific) and length was evaluated with the Fragment Analyzer Genomic DNA HS assay (Agilent). Samples were equimolar pooled in 8-plexes for SMRTbell library preparation and sequencing.

### FLAIRR-seq SMRTbell library preparation and sequencing

Eight-plex pools of targeted IgG or IgM amplicons were prepared into SMRTbell sequencing templates according to the “Procedure and Checklist for Iso-Seq Express Template for Sequel and Sequel II systems” protocol starting at the “DNA Damage Repair” step and using the SMRTbell Express Template Prep Kit 2.0, with some modifications (Pacific Biosciences). Briefly, targeted IgG and IgM amplicons underwent enzymatic DNA damage and end repair, followed by ligation with overhang SMRTbell adapters as specified in the protocol. To increase consistency in SMRTbell loading on the Sequel IIe system, we further treated the SMRTbell libraries with a nuclease cocktail to remove unligated amplified products, using the SMRTbell Enzyme Cleanup Kit, as recommended by the manufacturer (Pacific Biosciences). Briefly, after heat-killing the ligase with an incubation at 65°C, samples were treated with a nuclease cocktail at 37°C for 1 hour, and then purified with a 1.1X Pronex cleanup. Final SMRTbell libraries were evaluated for quantity and quality using the Qubit dsDNA HS assay and Fragment Analyzer Genomic DNA assay, respectively. Sequencing of each 8-plex, barcoded sample pool was performed on one SMRTcell 8M using primer v4 and polymerase v2.1 on the Sequel IIe system with 30 hr movies. Demultiplexed, high-fidelity circular consensus sequence reads (“HiFi reads”) were generated on the instrument for downstream analyses.

### AIRR-seq SMARTer Human BCR IgG/IgM sequencing

Matched healthy donor RNA was used to generate targeted IgG and IgM AIRR-seq libraries using the SMARTer Human BCR IgG IgM H/K/L Profiling Kit (Takara Bio USA) according to the manufacturer’s instructions with no modifications. Briefly, for each sample, proprietary IgG and IgM primers were used to amplify heavy chain transcripts following a 5’RACE reaction. AIRR-seq libraries were then quality controlled using the 2100 Bioanalyzer High Sensitivity DNA Assay Kit (Agilent) and the Qubit 3.0 Fluorometer dsDNA High Sensitivity Assay Kit. Sequencing on the MiSeq platform using 300 bp paired-end reads was performed using the 600-cycle MiSeq Reagent Kit v3 (Illumina) according to the manufacturer’s instructions, and FASTQ reads were generated using the associated DRAGEN software package (Illumina).

### B cell Iso-Seq

RNA extracted from healthy sorted B cells was used to generate Iso-Seq SMRTbell libraries following the “Procedure & Checklist Iso-Seq Express Template Preparation for the Sequel II System” with minor adaptations compared to the manufacturer’s instructions. Briefly, Iso-Seq libraries were generated using 500 ng high-quality (RIN > 8) RNA as input into oligo-dT primed cDNA synthesis (NEB). Barcoded primers were incorporated into the cDNA during second strand synthesis. Following double-stranded cDNA amplification, transcripts from two samples sourced from purified B cells and NK cells were equimolar pooled as previously described (33). SMRTbells were generated from the pooled cDNA as described above for the FLAIRR-seq amplification products, including the addition of a nuclease digestion step. Quantity and quality of the final Iso-Seq libraries were performed with the Qubit dsDNA High Sensitivity Assay Kit and Agilent Fragment Analyzer Genomic DNA assay, respectively. This 2-plex Iso-Seq pool was sequenced using primer v4 and polymerase v2.1 on the Sequel IIe system with a 30-hour movie. HiFi reads were generated on instrument before analyses. Demultiplexing of barcoded samples and generation of full-length non-concatemer (FLNC) predicted transcripts were performed using the Iso-Seq v3 pipeline available through SMRTLink (v.10.2). B-cell-derived FLNC reads were mapped to the human genome using the GMAP reference database and reads derived from chromosome 14 were extracted for downstream IGH transcript characterization via Immcantation, as described below.

### Immcantation analyses of IgG and IgM repertoires

Analyses of FLAIRR-seq, AIRR-seq, and Iso-seq datasets were performed using Immcantation tools (30, 31). Demultiplexed barcoded HiFi (for SMRT sequencing data) or FASTQ (for AIRR-seq) reads were first processed using the pRESTO tool for quality control, UMI processing, and error profiling (30). For AIRR-seq, pRESTO analysis data from paired-end reads (“R1” and “R2”) were trimmed to remove bases with < Q20 read quality and/or <125 bp length using the “FilterSeq trimqual” and “FilterSeq length”, respectively. IgG and IgM CH3 or CH4 primer sequences were identified with an error rate of 0.2, and primers identified were then noted in FASTQ headers using “MaskPrimers align”. Next, 12 basepair (bp) UMIs were located and extracted using “Maskprimers extract”. Sequences found to have the same UMIs were grouped and aligned using “AlignSets muscle,” with a consensus sequence generated for each UMI using “BuildConsensus”. Mate pairing of AIRR-seq reads was conducted using a reference-guided alignment requiring a minimum of a 5 bp overlap via “AssemblePairs sequential”. After collapsing consensus reads with the same UMI (“conscount”) using “CollapseSeq”,” reads with < 2 supporting sequences were removed from downstream analysis. For pRESTO processing of FLAIRR-seq, single HiFi reads (“R1”) reads did not require trimming due to > Q20 sequence quality across all bases. 5’TSO-UMI and CH3 or CH4 region primers were identified along with a 22 bp UMI with an error rate of 0.3 using “MaskPrimers align”. Reads were then grouped and aligned using “AlignSets muscle”. Due to the single molecule nature of FLAIRR-seq reads, no mate pairing was required. Consensus reads were then generated as described above, including removal of sequences with < 2 supporting reads. Read counts following each step of data filtration for AIRR-seq and FLAIRR-seq are represented in Supplementary Tables 3 and 4, respectively.

pRESTO-filtered reads for both AIRR-seq and FLAIRR-seq data were then input into the Change-O tool (Table 1). Iso-seq required no initial processing from pRESTO and was input into Change-O for IG gene reference alignment along with AIRR-seq and FLAIRR-seq data using “igblastn”, clonal clustering using “DefineClones”, and germline reconstruction and conversion using “CreateGermlines.py” and the GRCh38 chromosome 14 germline reference (31). Fully processed and annotated data was then converted into a TSV format for use in downstream analyses. The Alakazam Immcantation tool suite was then used to quantify gene usage analysis, calculate CDR3 length, assess somatic hypermutation frequencies, and analyze clonal diversity (31). SCOPer clonal assignment by spectral clustering was conducted for COVID-19 patient time course samples (34, 35). For clonal lineage tree analysis, the Dowser tool was used to examine clonal diversity and CSR over time (36).

**Table 1:** Comparative analysis metrics between AIRR-seq and FLAIRR-seq on matched samples

| Sample ID | Isotype | Method | RNA input (ng) | Input reads into Immcantation<sup>a</sup> | Post assembly /Single reads<sub>b</sub> | Unique VDJc | Unique Clones<sub>d</sub> | Unique CDR3<sub>e</sub> |
| --- | --- | --- | --- | --- | --- | --- | --- | --- |
| 0007 | IgG | AIRR | 100 | **841627** | 15217 | 3164 | 1008 | 1187 |
|  |  | FLAIRR | 335 | 181109 | 17881 | 4880 | 1358 | 1520 |
| 201c | IgG | AIRR | 100 | **1509063** | 25015 | 4568 | 1221 | 1371 |
|  |  | FLAIRR | 296 | 221301 | 36053 | 10819 | 2145 | 2388 |
| 203c | IgG | AIRR | 100 | **1006965** | 23085 | 5793 | 1436 | 1705 |
|  |  | FLAIRR | 106 | 475703 | 14923 | 3778 | 993 | 1138 |
| 602c | IgG | AIRR | 100 | **1275902** | 38078 | 7736 | 1542 | 2217 |
|  |  | FLAIRR | 149 | 344906 | 33903 | 9225 | 1828 | 2357 |
| 705c | IgG | AIRR | 100 | **1113660** | 15358 | 1362 | 557 | 576 |
|  |  | FLAIRR | 268 | 279601 | 18627 | 2328 | 1063 | 1081 |
| 1008 | IgG | AIRR | 100 | **1310350** | 56848 | 12217 | 3005 | 3307 |
|  |  | FLAIRR | 335 | 238820 | 57020 | 18235 | 3505 | 3813 |
| 1013 | IgG | AIRR | 100 | **1158222** | 25250 | 5027 | 1795 | 1964 |
|  |  | FLAIRR | 222 | 396322 | 27871 | 7487 | 2565 | 2744 |
| 2008 | IgG | AIRR | 100 | **835928** | 21681 | 3900 | 1235 | 1368 |
|  |  | FLAIRR | 240 | 418906 | 16441 | 4258 | 1308 | 1380 |
| 4002 | IgG | AIRR | 100 | **632272** | 8949 | 1112 | 727 | 772 |
|  |  | FLAIRR | 335 | 398838 | 19692 | 5377 | 1719 | 1887 |
| 5001 | IgG | AIRR | 100 | **695884** | 13930 | 1885 | 1134 | 1244 |
|  |  | FLAIRR | 335 | 313843 | 42845 | 12146 | 2226 | 2381 |
| 0007 | IgM | AIRR | 100 | **841627** | 12777 | 9154 | 7924 | 8163 |
|  |  | FLAIRR | 335 | 331380 | 46384 | 14926 | 12367 | 12851 |
| 201c | IgM | AIRR | 100 | **1509063** | 26850 | 17888 | 14281 | 14622 |
|  |  | FLAIRR | 296 | 148355 | 98069 | 12803 | 9353 | 9573 |
| 203c | IgM | AIRR | 100 | **1006965** | 24623 | 18189 | 16295 | 16629 |
|  |  | FLAIRR | 106 | 509536 | 37527 | 10636 | 9315 | 9419 |
| 602c | IgM | AIRR | 100 | **1275902** | 12306 | 8378 | 7704 | 7833 |
|  |  | FLAIRR | 149 | 273103 | 48897 | 16406 | 14495 | 14745 |
| 705c | IgM | AIRR | 100 | **1113660** | 19484 | 14273 | 12936 | 13413 |
|  |  | FLAIRR | 268 | 435391 | 79251 | 17417 | 15706 | 16249 |
| 1008 | IgM | AIRR | 100 | **1310350** | 20264 | 12710 | 7878 | 8368 |
|  |  | FLAIRR | 335 | 181063 | 47531 | 15284 | 8745 | 9276 |
| 1013 | IgM | AIRR | 100 | **1158222** | 12769 | 9012 | 8057 | 8241 |
|  |  | FLAIRR | 222 | 262617 | 52508 | 17812 | 15291 | 15759 |
| 2008 | IgM | AIRR | 100 | **835928** | 11608 | 8551 | 5838 | 6206 |
|  |  | FLAIRR | 240 | 502354 | 47596 | 14132 | 10225 | 10674 |
| 4002 | IgM | AIRR | 100 | **632272** | 18411 | 11607 | 11184 | 11367 |
|  |  | FLAIRR | 335 | 489287 | 91732 | 22379 | 19000 | 21532 |
| 5001 | IgM | FLAIRR | 335 | **313843** | 10018 | 12146 | 2226 | 2381 |
|  |  | FLAIRR | 355 | 171688 | 18944 | 6164 | 4866 | 5011 |

### Targeted IG gDNA capture, long-read sequencing, and IGenotyper analyses

FLAIRR-seq validation samples also underwent IGHC targeted enrichment and long-read sequencing as previously described (37). Briefly, gDNA was mechanically sheared and size selected to include 5-9kb fragments using the BluePippin (Sage Science). Samples were then end repaired and A-tailed using the standard KAPA library preparation protocol (Roche). Universal priming sequences and barcodes were then ligated onto samples for multiplexing (Pacific Biosciences). Barcoded gDNA libraries were captured using IGH-specific probes following a SeqCap protocol (Roche). 26 IGH-enriched samples were purified and pooled together for SMRTbell library prep as described above, including the final nuclease digestion step. Pooled SMRTbells were annealed to primer v4, bound to polymerase v2.0, and sequenced on the Sequel IIe system with 30h movies. After sequencing, HiFi reads were generated and analyzed by the IGenotyper pipeline (37). In brief, IGenotyper was used to detect single nucleotide variants and assemble sequences into haplotype-specific assemblies for downstream IGHC gene genotyping. Alleles were then extracted from assemblies using a bed file containing coordinates for each IGHC gene exon. After sequences were extracted, reads were then aligned to the IMGT database (downloaded on 2/21/22) and assigned as an exact match to IMGT, “novel” if there was no match to the IMGT database or “novel, extended” if a match was detected to a partial allele found in IMGT, but the IMGT allele was a substring of the IGenotyper identified allele (38). This set of alleles was then used as a ground truth dataset.

### IGHC gene genotyping with FLAIRR-seq

To genotype IGHC genes and alleles from FLAIRR-seq data, productive reads were filtered by IGHC length (900bp–1100bp) and aligned to the chromosome 14 hg38 reference using minimap2 along with SAMtools to generate sorted and indexed bam files (39, 40). WhatsHap was used to identify, genotype, and phase single nucleotide variants (SNV) (41). Phased SNVs were used to assign each read to a haplotype using MsPAC. Reads from each haplotype and gene were clustered using CD-HIT using a 100% identity clustering threshold parameter, and a single representative read from the largest cluster was aligned to the IGenotyper curated alleles using BLAST (42) to determine the closest matching IGHC gene and allele (38, 43). The representative read was selected based on 100% identity to all other sequences in that cluster.

### Inference of IGHV, IGHD, and IGHJ gene haplotypes from FLAIRR-seq data using IGHC gene anchors

To test the ability of IGHC genes to be used for the inference of IGHV, IGHD, and IGHJ haplotypes from FLAIRR-seq data, we chose one sample that was heterozygous for both IGHM and IGHJ6 (IGHJ6 is standardly used for AIRR-seq haplotype inference). For this sample, we employed TigGER (31, 44-46) to infer novel IGHV alleles, and generate sample-level IGHV genotypes using a Bayesian approach. Rearranged sequences within the Change-O table were then reannotated taking into account sample genotype and detected novel alleles. Updated annotations were then used to infer haplotypes using RAbHIT version 0.2.4 (47). Both IGHJ6 and IGHM were used as anchor points for haplotyping, and the resulting haplotypes were compared.

## Results

### Gene usage, CDR3, and SHM profiles characterized from FLAIRR-seq data are comparable to AIRR-seq and Iso-Seq

Current methods for commercially available 5’ RACE AIRR-Seq utilize targeted amplification of the variable region and, in some cases, a small portion of the first constant region exon (CH1), in conjunction with short-read sequencing to characterize IG repertoires. However, this minimal examination of the IGHC gene sequence is primarily used to define isotypes. No current method defines both the heavy chain variable and constant regions allowing for both subisotype classification and IGHC allele-level resolution. To address these technical limitations, we developed the FLAIRR-Seq method (Figure 1C), a targeted 5’ RACE approach combined with SMRT sequencing to generate highly accurate, near full-length IgG (∼1500 bp) and/or IgM (2000 bp) sequences, allowing for direct, simultaneous analysis of the heavy chain variable and constant regions (Figure 1D), including gene/allele identification for IGHV, IGHD, IGHJ and IGHC segments, and isotype- and subisotype-specific repertoire profiling.

To evaluate and validate the capabilities of FLAIRR-seq, matched FLAIRR-seq and AIRR-seq analyses were performed on ten healthy donor PBMC samples. FLAIRR-seq data were filtered from the initial HiFi reads (>Q20) to include only >Q40 reads. The average read quality of these filtered reads was >Q60 (99.9999%), with a pass filter rate ranging from 88%-93% of total reads. AIRR-seq FASTQ bases were trimmed to retain sequences with an average quality of Q20 (99%). These filtered reads were used as input into the Immcantation suite, specifically the pREST-O and Change-O tools, for IGHV, IGHD, and IGHJ gene assignment, and repertoire feature analyses, including identification of clones, extent of somatic hypermutation, and evaluation of CDR3 lengths. As shown in **Table 1**, fewer overall FLAIRR-seq reads were used as input into the Immcantation analyses, after required filtration and read assembly steps (which were not needed for the high-quality single-molecule FLAIRR-seq reads). FLAIRR-seq resulted in comparable or, in many cases, increased number of unique VDJ sequences, clones, and CDR3 sequences identified compared to the matched AIRR-seq-derived samples in both the IgG and IgM repertoires. These basic sequencing and initial analysis metrics demonstrated that FLAIRR-seq produced high-quality variable region data for detailed Ab repertoire analyses and is amenable to analysis using existing AIRR-seq analysis tools. While input RNA mass used for FLAIRR-seq was often more than for AIRR-seq, ongoing and future optimization of the method is aimed at reducing RNA input. Furthermore, a comparative cost analysis was performed. To obtain the closest metric to a direct comparison, we calculated the cost per “actionable read”, defined as the read number per sample and per method after read and length quality filtration and assembly, but prior to cluster consensus, performed in the pRESTO pipeline. This method was used to represent the total unique single molecule or assembled templates captured by either the FLAIRR-seq or AIRR-seq methods, respectively, that passed all necessary quality control criteria for downstream annotation and analyses irrespective of biologic repertoire diversity or clonality. Lastly, to remove the impact of pooling differences, we used this “per actionable read” cost to calculate the cost for the generation of 15,000 “actionable reads” as our standard price. For AIRR-seq this cost was $25.50 per sample, whereas for FLAIRR-seq, this cost was $33.57 per sample. These costs reflect needed reagents and consumables only, assume instrumentation access and do not include labor. Future optimization for FLAIRR-seq will include integrating a multiplexed array sequencing (MAS) step to concatenate reads and enhance overall depth of sequencing and multiplexing capacity per pool, resulting in decreasing costs per sample (48).

While optimizing FLAIRR-seq sample preparation, we examined whether upstream isolation of B cells before FLAIRR-seq molecular preparation would enhance the ability to detect IGHV, IGHD, and IGHJ gene usage. To do this, aliquots of PBMC (n=4) were split into two groups for RNA extraction: (i) RNA derived from bulk PBMC, and (ii) RNA isolated from purified pan B-cells, followed by FLAIRR-seq preparation, SMRT sequencing, and Immcantation analysis of both groups. IGHV, IGHD, and IGHJ gene usage correlations between groups are shown in Figure 2A and demonstrate a significant association (p-values ranging from 0.033 to 4.1e^-16^) strongly supporting the conclusion that B cell isolation before RNA extraction was not necessary to achieve comparable gene usage metrics. The limited differences that were observed could be explained by template sampling differences between the two experiments. Due to the strong associations observed and the ease of processing PBMC in bulk, we moved forward with RNA derived directly from PBMC aliquots for the remainder of our analyses.

**Figure 2.**
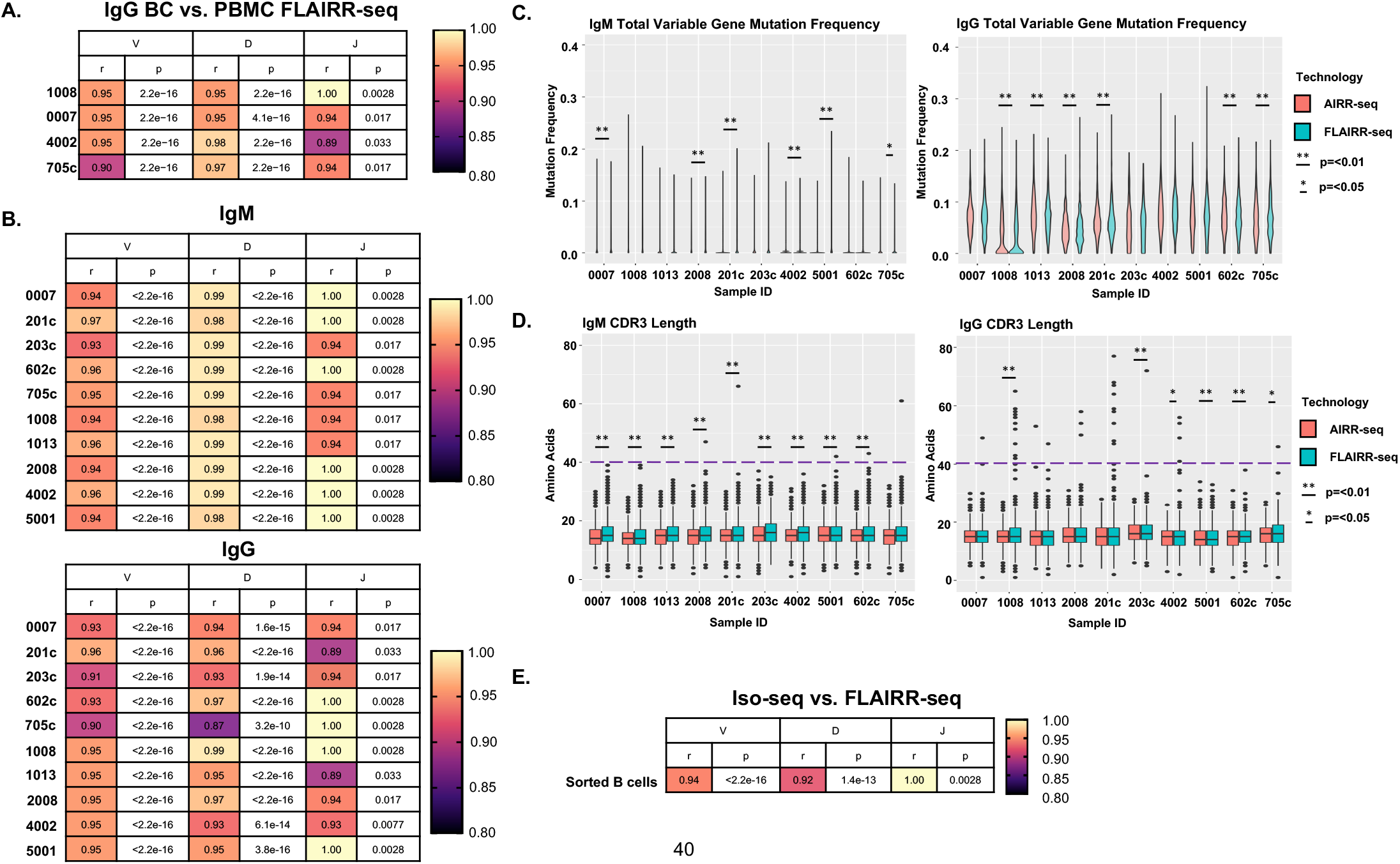
FLAIRR-seq shows robust characterization of V, D, and J genes. (A) Spearman ranked correlations and p-values of V, D, and J gene usage frequencies identified by FLAIRR-seq performed on matched total PBMC and purified B cells. (B) Heatmap of Spearman ranked correlations and p-values of V, D and J gene usage frequencies between FLAIRR-seq and AIRR-seq processed samples. (C) Boxplots of somatic hypermutation frequencies defined by FLAIRR-seq or AIRR-seq in IgM- and IgG-specific repertoires. (D) Boxplots of CDR3 length defined by FLAIRR-seq or AIRR-seq analysis in both IgM- and IgG-specific repertoires. (E) Spearman ranked correlation of V, D, and J gene usage frequencies between FLAIRR-seq-based and Iso-Seq-based repertoire profiling. Significant differences between FLAIRR-seq and AIRR-seq data indicated by * (p < 0.05) or ** (p < 0.01).

We established FLAIRR-seq performance by comparing its output to the commonly used 5’ RACE AIRR-seq method. 5’ RACE AIRR-seq was chosen as it provides resolution of the complete variable region and a small portion of IGHC, allowing for isotype differentiation. Following matched preparation of both FLAIRR-seq and AIRR-seq libraries from healthy donor PBMC samples (n=10), we compared multiple repertoire features to benchmark FLAIRR-seq performance. First, we evaluated IGHV, IGHD, and IGHJ gene usage frequencies. We observed significant correlations between FLAIRR-seq and AIRR-seq datasets in IGHV, IGHD and IGHJ gene usage for both IgM (V genes: r = 0.93-0.97, p=<2.2e^-16^; D genes: r = 0.98-0.99, p=<2.2e^-16^; J genes: r = 0.94-1.0, p = 0.0028-0.017) and IgG isotypes (V genes: r = 0.90-0.96, p=<2.2e^-16^; D genes: r = 0.87-0.99, p=<2.2e^-16^-6.1e^-14^;J genes: r = 0.89-1.0, p = 0.0028-0.033) (Figure 2B), indicating that FLAIRR-seq comparably resolves IGHV, IGHD, and IGHJ gene usage profiles. To note, IGHJ genes showed lower levels of significance (larger p-values) across all comparisons due to the relatively few genes that make up the IGHJ gene family compared to the more diverse IGHV and IGHD families. We next investigated the performance of both methods in terms of resolving somatic hypermutation (SHM) frequencies (Figure 2C), and complementarity determining region 3 (CDR3) lengths (Figure 2D), which are often used as measures of evaluating B cell affinity maturation. Although we did observe occasional statistically significant differences in the SHM frequency between AIRR-seq and FLAIRR-seq data using the same samples, these differences were not seen across all samples, suggesting that sample-to-sample variation may drive this observation rather than technology-based discrepancies. We found that CDR3 lengths were consistently longer in the FLAIRR-seq datasets for both the IgM and IgG isotypes in most donors. The characterization of unusually long CDR3 regions (> 40 nt) in the IgG sequences with FLAIRR-seq is likely due to the higher contiguity and quality afforded by the longer read lengths, which are less likely to be spanned by short-read 2×300 bp paired-end sequencing strategies. Together, these data demonstrate that FLAIRR-seq achieves comparable gene usage profiles, and improved resolution of long CDR3 sequences.

Others have recognized the power of long-read sequencing to resolve B cell repertoires using bulk Iso-Seq methods, allowing for the examination of full-length transcripts from isolated B cells (49). The Iso-Seq method captures full-length transcripts expressing a poly(A) tail without bias through oligo dT-based priming. The trade-offs of this approach are throughput, depth, and cost, as Iso-Seq processing generates a complete transcriptome per sample without enrichment of heavy chain sequences, which then need to be filtered out and analyzed, resulting in a considerable amount of non-repertoire data that is discarded. To investigate whether the untargeted transcriptome-wide Iso-Seq method would resolve a qualitatively different repertoire than FLAIRR-seq, which would have indicated FLAIRR-seq-driven primer bias, we performed matched Iso-Seq and FLAIRR-seq on purified B-cell derived RNA. IGHV, IGHD, and IGHJ gene usage frequencies were compared between Iso-Seq and FLAIRR-seq datasets (Spearman’s rank correlation), revealing significant correlations between usage profiles (V genes: r = 0.94, p=2.2e^-16^; D genes: r = 0.92, p =1.4e^-13^; J genes: r = 1.0, p = 0.0028; Figure 2E). These data strongly suggest that FLAIRR-seq has very limited to no primer-driven bias compared to whole transcriptome data. Collectively, this benchmarking dataset confirmed that FLAIRR-seq is comparable other state-of-the-art methods, providing robust characterization of commonly used repertoire metrics, with limited increases in per sample cost.

### IGenotyper and FLAIRR-seq provide constant region gene allele identification and allow for haplotyping of variable genes

The novel value added by FLAIRR-seq is improved resolution of IGHC, including estimation of IGHC gene and allele usage, subisotype identification, and phasing of variable and constant regions for comprehensive repertoire analysis. To evaluate the capabilities and accuracy of IGHC gene and allele identification with FLAIRR-seq, we first established a ground truth dataset of IGHC alleles for all 10 samples by targeted sequencing of the germline IGH locus (Figure 3A) using IGenotyper, as previously described (37). IGHG1, IGHG2, IGHG3, IGHG4 and IGHM alleles called by IGenotyper (see Methods) were assigned to one of three categories, schematized in Figure 3B: (i) “exact match” - alleles documented in the IMGT database; (ii) “novel not in IMGT” – alleles not documented in the IMGT database; or (iii) “extended” – alleles that matched partial alleles in the IMGT database (i.e., those only spanning a subset of exons), but were extended by sequences in our dataset. IGenotyper identified a total of 32 unique IGHG1, IGHG2, IGHG3, IGHG4 and IGHM alleles across all individuals, as schematized in Figure 3C. Among these 32 alleles, only 4 were documented in IMGT, the remaining represented novel alleles (n=11) or extensions (n=17) of known alleles. In aggregate, we observed a greater number of IGHG4 alleles than for any of the IGHG genes. Among these alleles were 4 sequences represented by suspected duplications of IGHG4. In fact, we observed 3 IGHG4 gene alleles in 4/10 samples, indicating the presence of gene duplications in these donors; in all cases, these alleles were also identified in the FLAIRR-seq data (see below). Given the relatively small size of this proof-of-concept healthy donor cohort, the identification of 28 (87%) novel or extended alleles underscores the extensive polymorphism In this region and reflects the paucity of information regarding this locus in existing immunogenomics databases.

**Figure 3.**
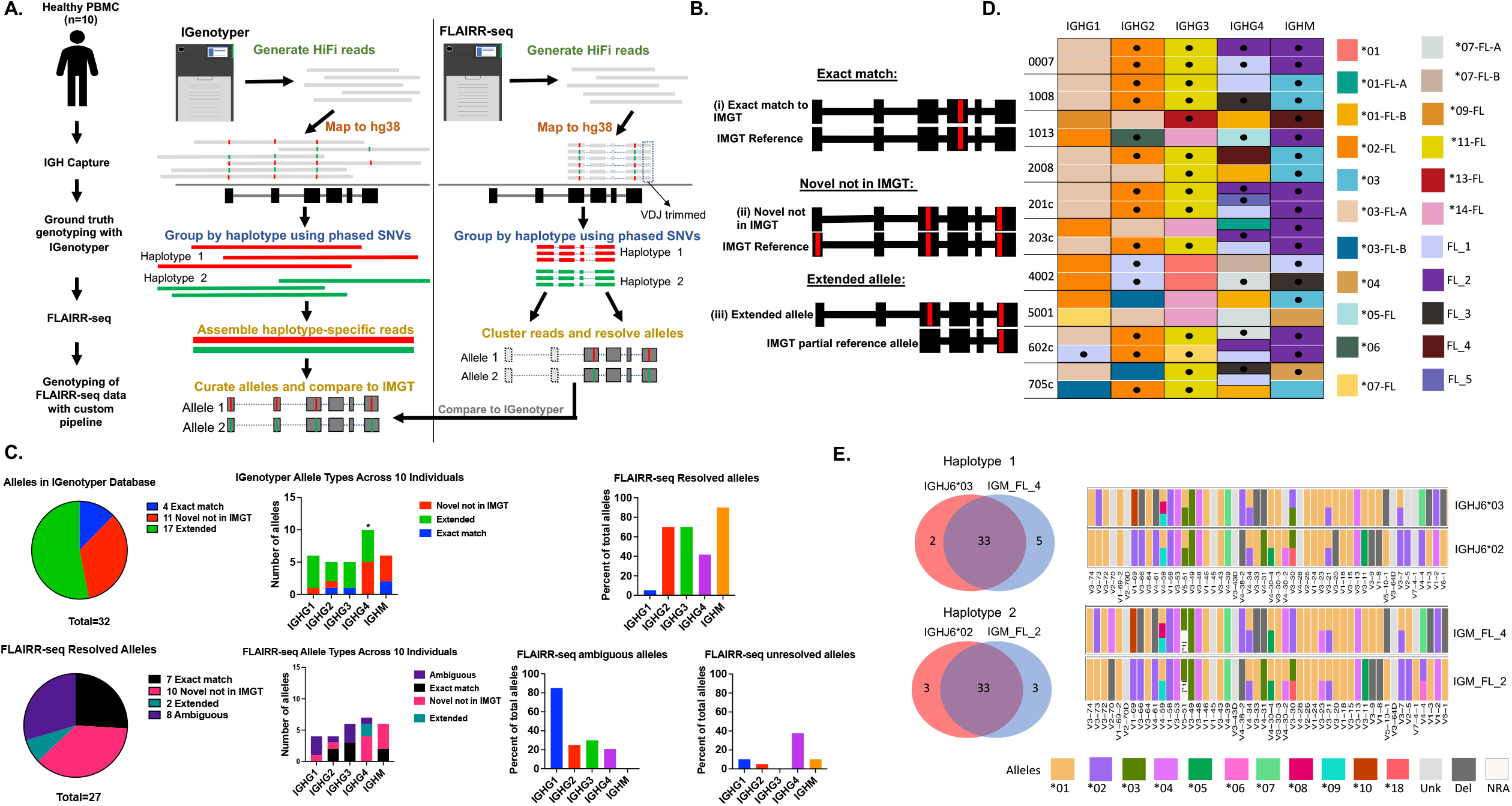
FLAIRR-seq provides novel IGHC resolution for allelic discovery and allows variable gene haplotyping. (A) Overview of experimental design and pipeline overviews of genotyping by IGenotyper (gDNA) and FLAIRR-seq (RNA). (B) Schematic depicting IGHC alleles identified by IGenotyper, partitioned by identification as (i) exact matches to documented IMGT alleles, (ii) novel alleles that are not in IMGT, or (iii) extended alleles. (C) Pie chart and stacked bar graph representing the total number of alleles and fraction of each category identified per IGHC gene as identified by either IGenotyper or FLAIRR-seq. Bar charts showing number of IGHC alleles from FLAIRR-seq that were resolved, ambiguous or unresolved when compared to IGenotyper alleles. * Indicates additional allele found due to IGHG4 duplication. (D) Table summarizing novel and extended alleles resolved by IGenotyper data. Extended alleles are denoted by *(allele number)-FL and novel alleles are denoted by FL_(number) alleles resolved by FLAIRR-seq are marked with a dot (•). (E) Venn diagrams showing number of IGHV haplotype allele/deletion calls when using IGHJ6 or IGHM anchors for each haplotype. Tile plots showing IGHV gene haplotypes inferred using either IGHJ6 anchors or IGHM anchors for one sample. Dark gray represents a deletion (DEL), off-white a non-reliable allele annotation (NRA), and light gray represents an unknown allele (Unk). Non-reliable alleles are annotated with an asterisk (*).

We next used the iGenotyper-derived IGHC gene database as the ground-truth for evaluating the capability of FLAIRR-seq to identify and resolve IGHC gene alleles. Based on our analysis workflow for identifying IGHC alleles from FLAIRR-seq data, we resolved 19/32 (59%) iGenotyper alleles at 100% identity; no additional false-positive alleles were identified. Of the alleles that were not unambiguously resolved by our FLAIRR-seq pipeline, 8 had allele defining single nucleotide variants (SNVs) 3’ of the FLAIRR-seq primers. The rate of true-positive allele calls using FLAIRR-seq across all 10 samples ranged from 5% for IGHG1 to 90% for IGHM (Figures 3C and 3D). As a result, the IGHC genotypes inferred by FLAIRR-seq have some limitations, but on the whole allow for much greater resolution of IGHC variation in the expressed repertoire than currently used methods. Future iterations of FLAIRR-seq will include primer optimization to facilitate better sequence coverage in the 3’ regions of the IGHC alleles and improve the direct genotyping capabilities of the FLAIRR-seq method.

Previous studies have demonstrated the use of IGHJ6 heterozygosity to infer haplotypes of V and D genes from AIRR-seq data (46, 50). However, the frequency of IGHJ6 heterozygotes in the population can vary. Therefore, we wanted to assess the utility of leveraging IGHC polymorphism resolved by FLAIRR-seq for haplotyping IGHV alleles with the publicly available tool RAbHIT(46, 50, 51). We selected a single donor (“1013”) from our cohort that was heterozygous for both IGHJ6 (*02 and *03) and IGHM (FL_2 and FL_4). Importantly, we were able to associate each IGHJ6 allele to the respective IGHM allele from the corresponding haplotype (Figure 3E). After defining germline IGHV alleles using TIgGER (31), we generated and compared IGHV haplotype inferences using either IGHJ6 or IGHM alleles as anchor genes using RAbHIT (47)(Figure 3E). Although some allele assignments were ambiguous (“unknown”) using both methods, we observed a strong consensus between haplotype inferences using the two anchor genes. For haplotype 1, represented by IGHJ6*03 and IGHM_FL_4, the IGHJ6*03-derived haplotype had 35 IGHV genes for which either an allele or deletion call was made. When using IGHM_FL_4, the same allele/deletion calls were made for 33 of these genes; in addition, using IGHM as the anchor gene, assignments were made for an additional 5 IGHV genes that had “unknown” designations using IGHJ6. Similarly, of the allele/deletion calls made for 36 IGHV genes on haplotype 2 assigned to IGHJ6*02, 33 of these gene had identical assignments to IGHM_FL_2. Together these results indicate that IGHC variants can be utilized for haplotype inference from repertoire data when commonly used IGHJ or IGHD genes are homozygous in individuals of interest.

### FLAIRR-seq enables isotype-, subisotype-, and allele-specific repertoire analyses

IGHG and IGHM alleles identified in each sample were used to annotate reads in each respective repertoire. These assignments allowed for partitioning of the repertoire by isotype, subisotype and IGHC allele (Figure 4). To demonstrate this, we utilized the same represtentative sample (“1013”) that was heterozygous for all IGHC genes. As shown in Figure 4A, IGHC gene assignments allow for subisotype and allele level frequencies to be estimated as a proportion of the overall IgG and IgM repertoires. In addition, detailed analyses of the repertoire can be conducted within each of these compartments. For example, Figure 4B shows the frequencies of IGHV gene subfamilies for each IGHG and IGHM allele identified in this sample. Using standard AIRR-seq analyses, we would not be able to identify allele-resolved V gene usage or enrichment within subisotype populations, which is important for linking subisotype functionality to particular antigen-specific VDJ clones.

**Figure 4.**
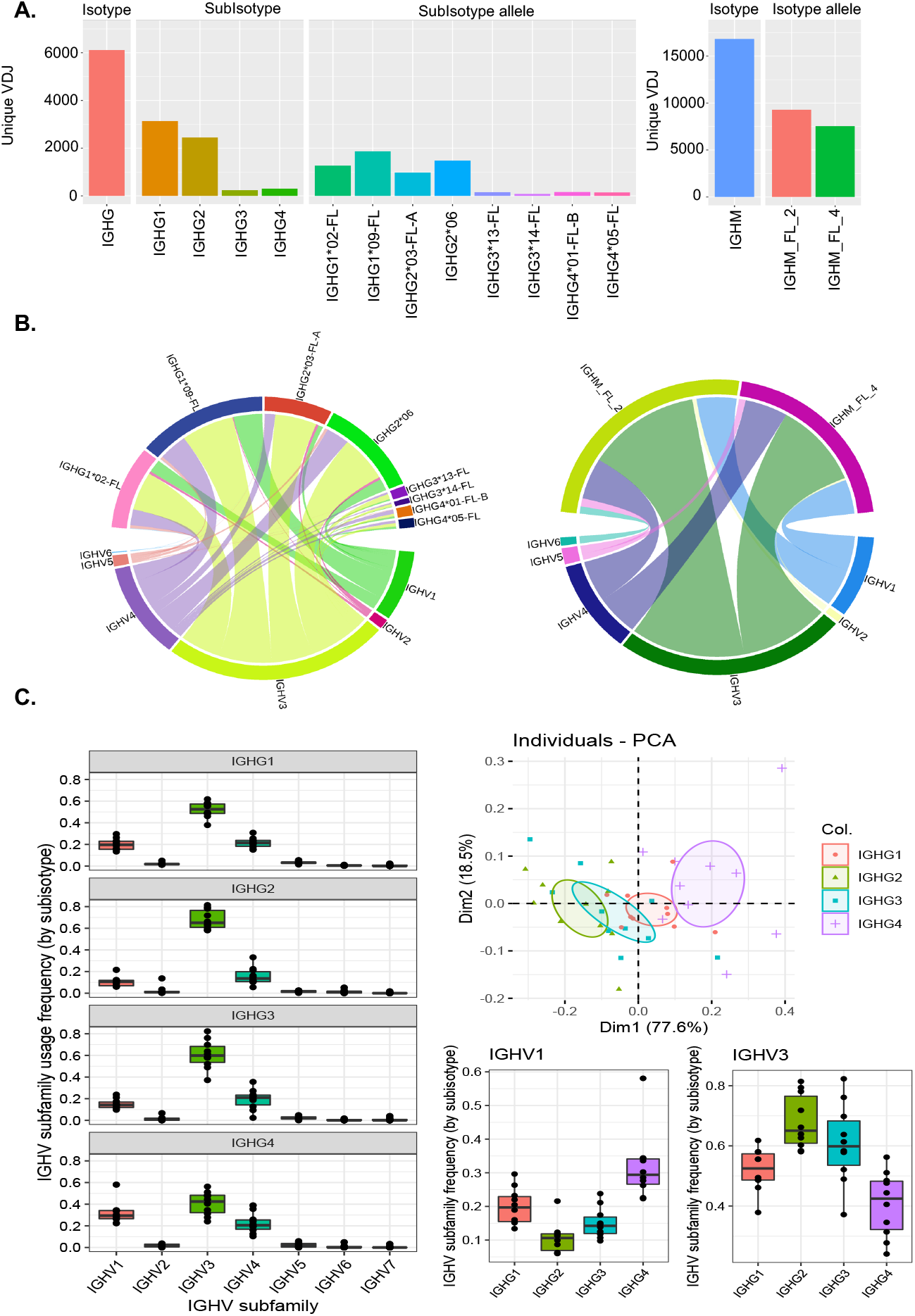
FLAIRR-seq resolves subisotype specific repertoire diversity. (A) Bar plots showing distribution of unique VDJ sequences across isotypes, subisotypes, and subisotype alleles in one representative sample, 1013, characterized by FLAIRR-seq. (B) Circos plots showing V family gene usage frequency within each subisotype allele for sample 1013. (C) Boxplots of V gene family usage frequencies within IGHG1, IGHG2, IGHG3, and IGHG4 repertoires across all ten individuals. (D) Principal component analysis of V gene family usage by subisotype; plot includes the first two principal components, and individual repertoires are colored by IGHG subisotype. (E) Boxplots showing sequence frequency of IGHV1 and IGHV3 family genes by subisotype across all 10 samples.

Through the partitioning of repertoire sequences by subisotype and IGHC allele, we found that FLAIRR-seq also allowed for trends to be assessed in aggregate across donors. To demonstrate this, we further examined V gene family usage partitioned by IgG subisotype across all 10 healthy donors. This analysis revealed expected patterns, in that IGHV1, IGHV3 and IGHV4 subfamily genes were dominant across the 4 subisotypes (Figure 4C). However, we did observe significant variation in subfamily proportions between subisotypes, associated with distinct profiles in specific subisotypes (Figure 4C). Specifically, the estimated frequencies of IGHV1 and IGHV3 were statistically different between subisotypes (P<0.01, ANOVA); IGHV1 usage was elevated in IGHG1 and IGHG4, whereas IGHV3 was elevated in IGHG2 and IGHG3. These analyses demonstrate the unique capability of FLAIRR-seq to examine variation in the expressed repertoire at the level of isotype, subisotype, and IGHC allele. As samples sizes increase, we expect that a multitude of additional repertoire features will become accessible to this kind of analysis leading to novel discoveries linking VDJ and IGHC genetic signatures.

### FLAIRR-seq identifies subisotype-specific clonal expansion and CSR in longitudinal samples

We wanted to investigate the utility of FLAIRR-seq in clinical samples, particularly to observe changes in immune repertoires over time. Ab responses are highly dynamic, with specific Ab clones expanding upon activation by antigen. We were interested to know if class switch recombination could be captured by FLAIRR-seq, given the capability to identify clones with the subisotype resolved. We had the opportunity to evaluate FLAIRR-seq resolved repertoires over time in four samples collected from one individual over their >13-day hospitalization for severe COVID-19 disease. Blood draws were taken on days 1, 4, 8, and 13 post-hospitalization (Figure 5A) and analyzed with FLAIRR-seq across all time points. After initial FLAIRR-seq processing and analysis, we defined unique clones using SCOPer, which clusters sequences based on CDR3 similarity and mutations in IGHV and IGHJ genes (34). This analysis allowed for the estimation of subisotype-specific clone counts across the four timepoints examined. Overall, we observed IgG1 dominated the repertoire at all four time points, but the proportion of subisotypes fluctuated over time (Figure 5B). Specifically, the IGHG2 and IGHG3-specific repertoires expanded from day 1 to day 4, but then contracted in overall frequency from day 8 to day 13 (Figure 5B). To assess clonal diversity within each subisotype repertoire across time, we calculated the Simpson’s diversity index (q=2) using Alakazam (31). All subisotype-specific repertoires became less diverse from day 1 to day 4, suggesting clonal expansion across the IGHG repertoire (Figure 5C). To note, IGHG4 was not included in diversity calculations because IGHG4 would have required higher sequencing depth to ascertain diversity, given the overall lower expression of IgG4 transcripts in this individual. Subisotype-specific repertoire polarity was also assessed by calculating the fraction of clones needed to represent 80% of the total repertoire (Figure 5D), with lower fractions representing more polarized and clonally expended repertoires. Results of this analysis were consistent with the diversity index, demonstrating an increase in clonal expansion (i.e., decreased polarity) at day 4 across IGHG1, IGHG2, and IGHG3 compartments, which returned to baseline at later timepoints.

**Figure 5.**
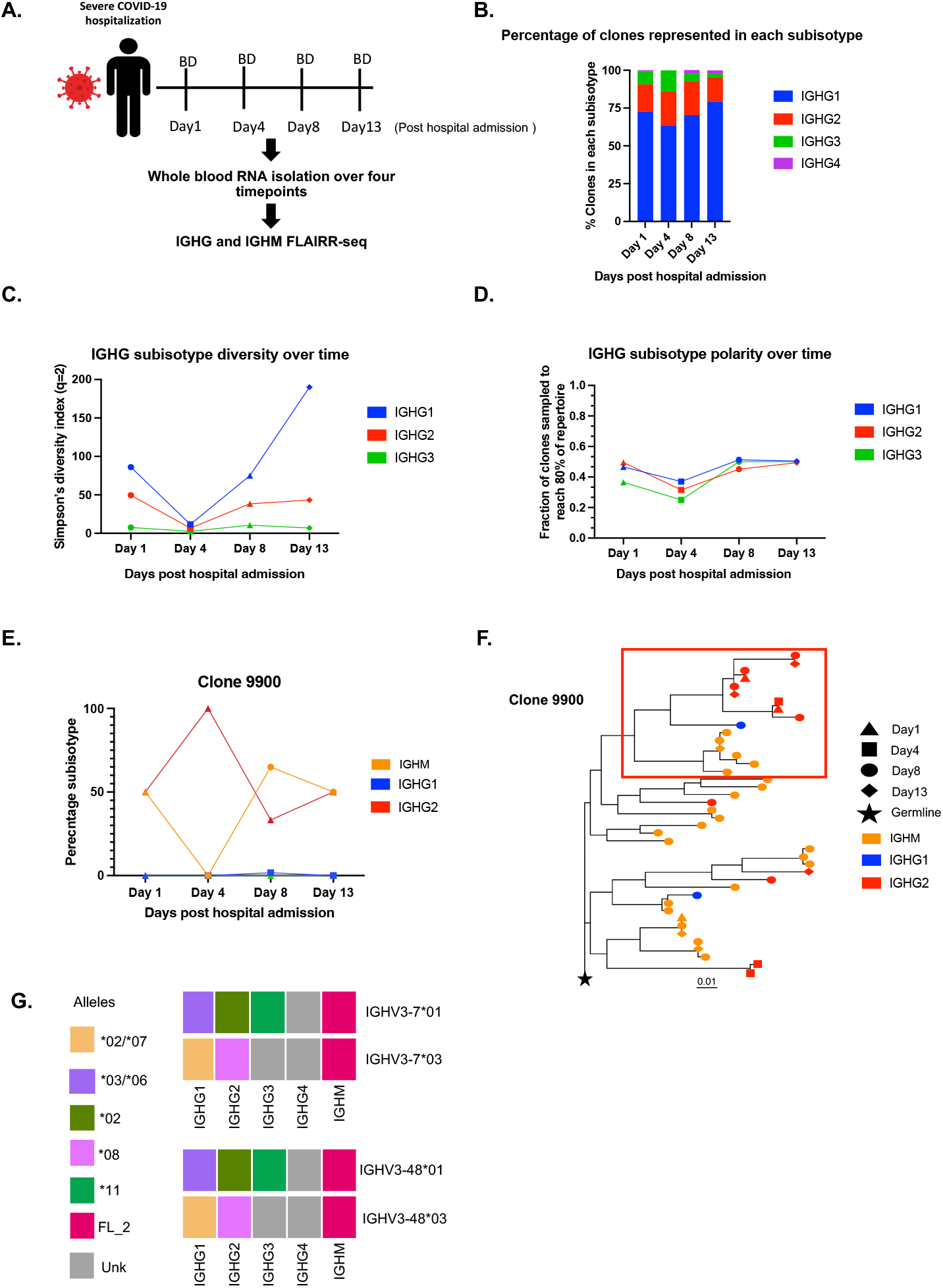
FLAIRR-seq resolves subisotype-specific clonal expansion and facilitates haplotype analysis of CSR in a patient hospitalized for COVID-19. (A) Overview of experimental design: whole blood-derived RNA was collected on days 1, 4, 8 and 13 post-hospitalization and used for FLAIRR-seq profiling. (B) Bar plot showing the percentage of clones represented by each subisotype across timepoints. (C) Simpson’s diversity index (q=2) for all clones in each subisotype across four timepoints; IgG4 not included due to low sequence counts. (D) Polarity, or the fraction of clones needed to comprise 80% of the repertoire, reported as fraction of total subisotype-specific repertoires across time. (E) Distribution of a single clone “9900” across isotypes and subisotypes over time, suggesting CSR of this clone. (F) Phylogenetic tree constructed from sequences/members of clone 9900, with the inferred germline sequence as the outgroup (star). Shapes and colors of tips (sequences) indicate time point and isotype/subisotype. The scale bar represents the number of mutations between each node in the tree. The subclade within the red box is represented by multiple time points and subisotypes, providing evidence of CSR. (G) Tile plot showing the assignment of IGHC alleles to their respective haplotypes, based on the frequency of observations in which each IGHC allele was linked to each respective allele of heterozygous V genes, IGHV3-7 and IGHV3-48; light gray denotes IGHC alleles for which haplotype assignment was not possible. Analysis of sequences in (F) revealed that the IGHG1 and IGHG2 alleles represented in the phylogeny came from the same haplotype (IGHG1*02/*07, IGHG2*08, IGHM_FL_2).

CSR mediates the switching of Abs from one sub/isotype to another. This occurs through the somatic recombination of IGHC genes, which brings the switched/selected IGHC genes adjacent to the recombined IGHV, IGHD, and IGHJ segments, facilitating transcription. The switching of isotypes and subisotypes can result in changes to associated effector functions of the Ab while maintaining antigen-specific variable regions. Given the ability of FLAIRR-seq to resolve clones with subisotype and IGHC allele resolution, as proof-of-concept, we sought to assess whether FLAIRR-seq could allow for more detailed haplotype-level analysis of CSR through the course of infection. To do this, we identified the largest clones in our dataset that were both represented by multiple isotypes and found across timepoints. In total, using SCOper (34), we identified 19 unique clonal lineages that met this criteria. We focused our detailed analysis on one of the largest clones, “9900” (Figure 5E), comprising IGHM, IGHG1, and IGHG2 sequences. On day 1 post-hospitalization, clone 9900 sequences were represented by both IGHM and IGHG2. At day 4, IGHG2 was the only isotype observed, whereas again at day 8, both IGHM and IGHG2 were observed, as well as IGHG1. To visualize CSR, we built a phylogeny using Dowser (36). Highlighted in the red box on the phylogenetic tree shown in Figure 5F, we observe a single subclade that is represented by IGHM, IGHG1, and IGHG2.

We were also able to resolve IGHC alleles from this individual, with the exception of IGHG1 alleles which were ambiguous. Critically, both IGHG1 and IGHG2 were heterozygous (Figure 5G). Through the assignment IGHG alleles to haplotypes within this individual using heterozygous V genes (IGHV3-7 and IGHV3-48), we were able to determine that sequences from clone 9900 (Figure 5F) utilized IGHG1 and IGHG2 alleles from the same haplotype, associated with IGHV3-7*07 and IGHV3-48*03 (31). This observation offers direct characterization of CSR events occurring on the same chromosome. When we looked across the remaining clones (n=8) in this dataset that spanned time points and were represented by IGHG1 and IGHG2 subisotypes, we were able to confirm that all of these clones used IGHG alleles from the same haplotype.

Together, these data provide demonstrative proof-of-concept evidence that FLAIRR-seq profiling performs robustly on clinical samples, including RNA directly extracted from whole blood. In addition, these data provide novel repertoire resolution extending what would have been possible with standard AIRR-seq methods, including analysis of subisotype-specific repertories, evaluation of clonal expansion, and characterization of CSR in the IgG and IgM repertoires.

## Discussion

Here we present the development, validation, and application of FLAIRR-seq, a novel method to resolve near full-length Ab transcripts from bulk PBMC-, isolated B cell- and whole blood-derived total RNA. FLAIRR-seq enables highly accurate, simultaneous resolution of variable and constant regions and suggests that IGHC polymorphism is far more extensive than previously assumed. FLAIRR-seq performed equivalent to or with increased resolution compared to existing standard 5’RACE AIRR-seq methods when resolving V, D, and J gene calls, CDR3 lengths, and SHM signatures, suggesting that our CH3/CH4 targeting strategies did not compromise variable region characterization while simultaneously adding the capability to resolve IGHC variation. Little to no primer bias was observed when compared to Ab repertoire profiling from total mRNA Iso-seq methods. FLAIRR-seq provides the novel ability to use IGHC gene usage to identify subisotypes and genotype heavy chain transcripts, linking these data back to evaluate subisotype-specific repertoires, clonal expansion, and CSR. Underscoring the underappreciated extent of IGHC variation, our profiling of a restricted cohort of only 10 individuals from relatively homogenous backgrounds still identified 4 and 7 completely novel IGHC alleles in IgM and IgG, respectively, and extended an additional 17 alleles beyond which had been available in the IMGT database.

The unique capabilities of FLAIRR-seq will allow for novel examination of Ab repertoires, including the characterization of variable gene usage and clonotype distribution within unique subisotype subsets. This perspective has the potential to provide key insights into dynamic Ab responses in diseases known to be mediated by subisotype-specific processes or have skewed subisotype distribution as predictive markers of disease, including Myasthenia gravis (mediated by pathogenic autoantibodies across subisotypes that give rise to varied disease pathologies), Acute Rheumatic Fever (associated with elevated IgG3), and melanoma (skewing towards IgG4 in late-stage disease thought to be indicative of tolerogenic responses and poor prognosis)(12-14). These subisotype-specific repertoire profiling approaches may be the first step toward identification of unique clones that mediate disease pathogenicity or serve as high-resolution biomarkers to disease progression, as well as open the door for potential functional experiments on subisotype clones of interest, including examining the functional impact of the novel IGHC alleles identified here. Expanded population-based FLAIRR-seq profiling and curation of novel IGHC alleles, particularly in conjunction with IGenotyper targeted genomic assembly efforts in IGH, will be a significant first step in defining the full extent of variation in a region too long assumed to be relatively invariant.

The Fc region is known to be critical for modulating differential Ab effector functions. These differential functionalities are currently understood to be regulated by differential posttranslational modifications, such as variable glycosylation (52-55). Future FLAIRR-seq profiling will be a valuable tool to investigate how genomic variation across IGHC genes impacts residue usage and resultant Fc receptor binding, signaling potential, crosslinking, and potential for posttranslational modification, all of which would be expected to alter downstream effector functions, such as ADCC, ADCP, and complement fixation.

We further demonstrate that FLAIRR-seq can effectively examine clonal expansion and CSR in longitudinal samples, demonstrating the feasibility of using FLAIRR-seq to resolve Ab repertoire dynamics. This increased resolution will further our understanding of Ab repertoire evolution in the transition of acute to chronic disease states, many of which are associated with overall IgG subisotype distribution changes that are thought to reflect the inflammatory milieu (14, 56). One example is advanced melanoma, where late-stage disease is characterized by elevated IgG4 compared to IgG1, which is believed to reflect a more tolerizing, pro-tumor environment (12). FLAIRR-seq examination of these samples may identify specific repertoire distribution patterns that could act as biomarkers of disease progression. Moving forward it is critical to account for all variability within the Ab repertoire for the most comprehensive understanding of repertoire dynamics and the myriad factors impacting Ab effector function. Future efforts will expand FLAIRR-seq methods to target IGHA and IGHE repertoires, as well as implement multiplexed arrays (MAS) sequencing to considerably increase throughput and lower cost (48). Together, the data presented here demonstrate that the FLAIRR-seq method provides a comprehensive characterization of allele-resolved IgG and IgM repertoires, detailing variable region gene usage and measurements of maturation, isotype and subisotype identification, and the unappreciated extent of constant region variation, which will be necessary to fully appreciate the impact of IG genomic variation in health and disease.

## Acknowledgements

The authors would like to thank Kamille Rasche, Kaitlyin Shields, Uddalok Jana, and the staff of the University of Louisville Sequencing Technology Center for assistance in the sequencing and analysis of FLAIRR-seq samples. They would also like to acknowledge the support and assistance from Gur Yaari and Ayelet Peres for their assistance with the RabHIT workflow. This work was funded, in part, by P20 GM135004-02 (EEF, DT, CTW, and MLS).

## Notes

### Competing Interest Statement

The authors have declared no competing interest.

### Summary of Updates

Add contributing authors.

## References

1. Schroeder, H. W., and L. Cavacini. 2010. Structure and function of immunoglobulins. Journal of Allergy and Clinical Immunology 125: S41–S52.

2. Janda, A., A. Bowen, N. S. Greenspan, and A. Casadevall. 2016. Ig Constant Region Effects on Variable Region Structure and Function. Front Microbiol 7: 22.

3. Nakano, T., M. Matsui, I. Inoue, T. Awata, S. Katayama, and T. Murakoshi. 2011. Free immunoglobulin light chain: Its biology and implications in diseases. Clinica Chimica Acta 412: 843–849.

4. Lu, L. L., T. J. Suscovich, S. M. Fortune, and G. Alter. 2018. Beyond binding: antibody effector functions in infectious diseases. Nature Reviews Immunology 18: 46–61.

5. Vidarsson, G., G. Dekkers, and T. Rispens. 2014. IgG subclasses and allotypes: from structure to effector functions. Front Immunol 5: 520–520.

6. Greiff, V., C. R. Weber, J. Palme, U. Bodenhofer, E. Miho, U. Menzel, and S. T. Reddy. 2017. Learning the High-Dimensional Immunogenomic Features That Predict Public and Private Antibody Repertoires. The Journal of Immunology 199: 2985.

7. Tonegawa, S. 1983. Somatic generation of antibody diversity. Nature 302: 575–581.

8. Nishana, M., and S. C. Raghavan. 2012. Role of recombination activating genes in the generation of antigen receptor diversity and beyond. Immunology 137: 271–281.

9. Tong, P., A. Granato, T. Zuo, N. Chaudhary, A. Zuiani, S. S. Han, R. Donthula, A. Shrestha, D. Sen, J. M. Magee, M. P. Gallagher, C. E. van der Poel, M. C. Carroll, and D. R. Wesemann. 2017. IgH isotype-specific B cell receptor expression influences B cell fate. Proc Natl Acad Sci U S A 114: E8411–e8420.

10. Noviski, M., J. L. Mueller, A. Satterthwaite, L. A. Garrett-Sinha, F. Brombacher, and J. Zikherman. 2018. IgM and IgD B cell receptors differentially respond to endogenous antigens and control B cell fate. eLife 7: e35074.

11. Stavnezer, J., J. E. Guikema, and C. E. Schrader. 2008. Mechanism and regulation of class switch recombination. Annu Rev Immunol 26: 261–292.

12. Karagiannis, P., A. E. Gilbert, D. H. Josephs, N. Ali, T. Dodev, L. Saul, I. Correa, L. Roberts, E. Beddowes, A. Koers, C. Hobbs, S. Ferreira, J. L. Geh, C. Healy, M. Harries, K. M. Acland, P. J. Blower, T. Mitchell, D. J. Fear, J. F. Spicer, K. E. Lacy, F. O. Nestle, and S. N. Karagiannis. 2013. IgG4 subclass antibodies impair antitumor immunity in melanoma. J Clin Invest 123: 1457–1474.

13. Chung, A. W., T. K. Ho, P. Hanson-Manful, S. Tritscheller, J. M. Raynes, A. L. Whitcombe, M. L. Tay, R. McGregor, N. Lorenz, J. R. Oliver, J. K. Gurney, C. G. Print, N. J. Wilson, W. J. Martin, D. A. Williamson, M. G. Baker, and N. J. Moreland. 2020. Systems immunology reveals a linked IgG3-C4 response in patients with acute rheumatic fever. Immunol Cell Biol 98: 12–21.

14. Vander Heiden, J. A., P. Stathopoulos, J. Q. Zhou, L. Chen, T. J. Gilbert, C. R. Bolen, R. J. Barohn, M. M. Dimachkie, E. Ciafaloni, T. J. Broering, F. Vigneault, R. J. Nowak, S. H. Kleinstein, and K. C. O’Connor. 2017. Dysregulation of B Cell Repertoire Formation in Myasthenia Gravis Patients Revealed through Deep Sequencing. J Immunol 198: 1460–1473.

15. Huijbers, M. G., L. A. Querol, E. H. Niks, J. J. Plomp, S. M. van der Maarel, F. Graus, J. Dalmau, I. Illa, and J. J. Verschuuren. 2015. The expanding field of IgG4-mediated neurological autoimmune disorders. European Journal of Neurology 22: 1151–1161.

16. Polonelli, L., J. Pontón, N. Elguezabal, M. D. Moragues, C. Casoli, E. Pilotti, P. Ronzi, A. S. Dobroff, E. G. Rodrigues, M. A. Juliano, D. L. Maffei, W. Magliani, S. Conti, and L. R. Travassos. 2008. Antibody complementarity-determining regions (CDRs) can display differential antimicrobial, antiviral and antitumor activities. PLoS One 3: e2371.

17. Liu, H., W. Pan, C. Tang, Y. Tang, H. Wu, A. Yoshimura, Y. Deng, N. He, and S. Li. 2021. The methods and advances of adaptive immune receptors repertoire sequencing. Theranostics 11: 8945–8963.

18. Trück, J., A. Eugster, P. Barennes, C. M. Tipton, E. T. Luning Prak, D. Bagnara, C. Soto, J. S. Sherkow, A. S. Payne, M.-P. Lefranc, A. Farmer, A. C. The, M. Bostick, and E. Mariotti-Ferrandiz. 2021. Biological controls for standardization and interpretation of adaptive immune receptor repertoire profiling. eLife 10: e66274.

19. Horns, F., C. Vollmers, D. Croote, S. F. Mackey, G. E. Swan, C. L. Dekker, M. M. Davis, and S. R. Quake. 2016. Lineage tracing of human B cells reveals the in vivo landscape of human antibody class switching. eLife 5: e16578.

20. Calonga-Solís, V., D. Malheiros, M. H. Beltrame, L. d. B. Vargas, R. M. Dourado, H. C. Issler, R. Wassem, M. L. Petzl-Erler, and D. G. Augusto. 2019. Unveiling the Diversity of Immunoglobulin Heavy Constant Gamma (IGHG) Gene Segments in Brazilian Populations Reveals 28 Novel Alleles and Evidence of Gene Conversion and Natural Selection. Front Immunol 10.

21. Jonsson, S., G. Sveinbjornsson, A. L. de Lapuente Portilla, B. Swaminathan, R. Plomp, G. Dekkers, R. Ajore, M. Ali, A. E. H. Bentlage, E. Elmér, G. I. Eyjolfsson, S. A. Gudjonsson, U. Gullberg, A. Gylfason, B. V. Halldorsson, M. Hansson, H. Holm, Å. Johansson, E. Johnsson, A. Jonasdottir, B. R. Ludviksson, A. Oddsson, I. Olafsson, S. Olafsson, O. Sigurdardottir, A. Sigurdsson, L. Stefansdottir, G. Masson, P. Sulem, M. Wuhrer, A.-K. Wihlborg, G. Thorleifsson, D. F. Gudbjartsson, U. Thorsteinsdottir, G. Vidarsson, I. Jonsdottir, B. Nilsson, and K. Stefansson. 2017. Identification of sequence variants influencing immunoglobulin levels. Nature Genetics 49: 1182–1191.

22. Buck, D., E. Albrecht, M. Aslam, A. Goris, N. Hauenstein, A. Jochim, S. Cepok, V. Grummel, B. Dubois, A. Berthele, P. Lichtner, C. Gieger, J. Winkelmann, and B. Hemmer. 2013. Genetic variants in the immunoglobulin heavy chain locus are associated with the IgG index in multiple sclerosis. Ann Neurol 73: 86–94.

23. Keyeux, G., G. Lefranc, and M.-P. Lefranc. 1989. A multigene deletion in the human IGH constant region locus involves highly homologous hot spots of recombination. Genomics 5: 431–441.

24. Bashirova, A. A., W. Zheng, M. Akdag, D. G. Augusto, N. Vince, K. L. Dong, C. O’HUigin, and M. Carrington. 2021. Population-specific diversity of the immunoglobulin constant heavy G chain (IGHG) genes. Genes Immun 22: 327–334.

25. Lefranc, M. P., G. Lefranc, G. de Lange, T. A. Out, P. J. van den Broek, J. van Nieuwkoop, J. Radl, A. N. Helal, H. Chaabani, E. van Loghem, and et al. 1983. Instability of the human immunoglobulin heavy chain constant region locus indicated by different inherited chromosomal deletions. Mol Biol Med 1: 207–217.

26. Lefranc, M.-P., and G. Lefranc. 2012. Human Gm, Km, and Am Allotypes and Their Molecular Characterization: A Remarkable Demonstration of Polymorphism. In Immunogenetics: Methods and Applications in Clinical Practice. F. T. Christiansen, and B. D. Tait, eds. Humana Press, Totowa, NJ. 635–680.

27. Lefranc, M.-P., G. Lefranc, and T. H. Rabbitts. 1982. Inherited deletion of immunoglobulin heavy chain constant region genes in normal human individuals. Nature 300: 760–762.

28. van Erp, E. A., W. Luytjes, G. Ferwerda, and P. B. van Kasteren. 2019. Fc-Mediated Antibody Effector Functions During Respiratory Syncytial Virus Infection and Disease. Front Immunol 10.

29. Jefferis, R., J. Lund, and J. D. Pound. 1998. IgG-Fc-mediated effector functions: molecular definition of interaction sites for effector ligands and the role of glycosylation. Immunological Reviews 163: 59–76.

30. Vander Heiden, J. A., G. Yaari, M. Uduman, J. N. H. Stern, K. C. O’Connor, D. A. Hafler, F. Vigneault, and S. H. Kleinstein. 2014. pRESTO: a toolkit for processing high-throughput sequencing raw reads of lymphocyte receptor repertoires. Bioinformatics 30: 1930–1932.

31. Gupta, N. T., J. A. Vander Heiden, M. Uduman, D. Gadala-Maria, G. Yaari, and S. H. Kleinstein. 2015. Change-O: a toolkit for analyzing large-scale B cell immunoglobulin repertoire sequencing data. Bioinformatics 31: 3356–3358.

32. Charney, A. W., N. W. Simons, K. Mouskas, L. Lepow, E. Cheng, J. Le Berichel, C. Chang, R. Marvin, D. M. Del Valle, S. Calorossi, A. Lansky, L. Walker, M. Patel, H. Xie, N. Yi, A. Yu, G. Kang, A. Mendoza, L. E. Liharska, E. Moya, M. Hartnett, S. Hatem, L. Wilkins, M. Eaton, H. Jamal, K. Tuballes, S. T. Chen, A. Tabachnikova, J. Chung, J. Harris, C. Batchelor, J. Lacunza, M. Yishak, K. Argueta, N. Karekar, B. Lee, G. Kelly, D. Geanon, D. Handler, J. Leech, H. Stefanos, T. Dawson, I. Scott, N. Francoeur, J. S. Johnson, A. Vaid, B. S. Glicksberg, G. N. Nadkarni, E. E. Schadt, B. D. Gelb, A. Rahman, R. Sebra, G. Martin, C. Agashe, P. Agrawal, A. Akyatan, K. Alesso-Carra, E. Alibo, K. Alvarez, A. Amabile, S. Ascolillo, R. Bailey, P. Begani, P. B. Correra, S.-A. Brown, M. Buckup, L. Burka, L. Cambron, G. Carrara, S. Chang, J. Chien, M. Chowdhury, C. C. Bozkus, P. Comella, D. Cosgrove, F. Cossarini, L. Cotter, A. Dave, B. Dayal, M. Dhainaut, R. Dornfeld, K. Dul, N. Eber, C. Elaiho, F. Fabris, J. Faith, D. Falci, S. Feng, B. Fennessy, M. Fernandes, S. Gangadharan, J. Grabowska, G. Gyimesi, M. Hamdani, M. Herbinet, E. Herrera, A. Hochman, G. E. Hoffman, J. Hook, L. Horta, E. Humblin, S. Karim, J. Kim, D. Lebovitch, G. Lee, G. H. Lee, J. Lee, M. Leventhal, K. Lindblad, A. Livanos, R. Machado, Z. Mahmood, K. Mar, S. Maskey, P. Matthews, K. Meckel, S. Mehandru, C. Mercedes, D. Meyer, G. Mollaoglu, S. Morris, K. Nie, M. Nisenholtz, G. Ofori-Amanfo, K. Onel, M. Ounadjela, V. Patel, C. Pruitt, S. Rathi, J. Redes, I. Reyes-Torres, A. Rodrigues, A. Rodriguez, V. Roudko, E. Ruiz, P. Scalzo, P. Silva, A. S. Schanoski, M. Straw, S. Tabachnikova, C. Teague, B. Upadhyaya, V. Van Der Heide, N. Vaninov, D. Wacker, H. Walsh, C. M. Wilk, J. Wilson, K. M. Wilson, L. Xue, N.-a. Yeboah, S. Young, N. Zaks, R. Zha, T. Marron, N. Beckmann, S. Kim-Schulze, S. Gnjatic, M. Merad, and C.-B. T. The Mount Sinai. 2020. Sampling the host response to SARS-CoV-2 in hospitals under siege. Nature Medicine 26: 1157–1158.

33. Woolley, C., J. Chariker, E. Rouchka, E. Ford, E. Hudson, S. Waigel, M. Smith, and T. Mitchell. 2022. Reference long-read isoform-aware transcriptomes of four human peripheral blood lymphocyte subsets G3: Genes, Genomes, Genetics

34. Nouri, N., and S. H. Kleinstein. 2018. A spectral clustering-based method for identifying clones from high-throughput B cell repertoire sequencing data. Bioinformatics 34: i341–i349.

35. Nouri, N., and S. H. Kleinstein. 2020. Somatic hypermutation analysis for improved identification of B cell clonal families from next-generation sequencing data. PLOS Computational Biology 16: e1007977.

36. Hoehn, K. B., O. G. Pybus, and S. H. Kleinstein. 2022. Phylogenetic analysis of migration, differentiation, and class switching in B cells. PLOS Computational Biology 18: e1009885.

37. Rodriguez, O. L., W. S. Gibson, T. Parks, M. Emery, J. Powell, M. Strahl, G. Deikus, K. Auckland, E. E. Eichler, W. A. Marasco, R. Sebra, A. J. Sharp, M. L. Smith, A. Bashir, and C. T. Watson. 2020. A Novel Framework for Characterizing Genomic Haplotype Diversity in the Human Immunoglobulin Heavy Chain Locus. Front Immunol 11.

38. Lefranc, M. P. 2001. IMGT, the international ImMunoGeneTics database. Nucleic Acids Res 29: 207–209.

39. Li, H. 2018. Minimap2: pairwise alignment for nucleotide sequences. Bioinformatics 34: 3094–3100.

40. Li, H., B. Handsaker, A. Wysoker, T. Fennell, J. Ruan, N. Homer, G. Marth, G. Abecasis, R. Durbin, and G. P. D. P. Subgroup. 2009. The Sequence Alignment/Map format and SAMtools. Bioinformatics 25: 2078–2079.

41. Martin, M., M. Patterson, S. Garg, S. O Fischer, N. Pisanti, G. W. Klau, A. Schöenhuth, and T. Marschall. 2016. WhatsHap: fast and accurate read-based phasing. bioRxiv: 085050.

42. Altschul, S. F., W. Gish, W. Miller, E. W. Myers, and D. J. Lipman. 1990. Basic local alignment search tool. J Mol Biol 215: 403–410.

43. Fu, L., B. Niu, Z. Zhu, S. Wu, and W. Li. 2012. CD-HIT: accelerated for clustering the next-generation sequencing data. Bioinformatics (Oxford, England) 28: 3150–3152.

44. Gadala-Maria, D., G. Yaari, M. Uduman, and S. H. Kleinstein. 2015. Automated analysis of high-throughput B-cell sequencing data reveals a high frequency of novel immunoglobulin V gene segment alleles. Proceedings of the National Academy of Sciences 112: E862–E870.

45. Gadala-Maria, D., M. Gidoni, S. Marquez, J. A. Vander Heiden, J. T. Kos, C. T. Watson, K. C. O’Connor, G. Yaari, and S. H. Kleinstein. 2019. Identification of Subject-Specific Immunoglobulin Alleles From Expressed Repertoire Sequencing Data. Front Immunol 10.

46. Gidoni, M., O. Snir, A. Peres, P. Polak, I. Lindeman, I. Mikocziova, V. K. Sarna, K. E. A. Lundin, C. Clouser, F. Vigneault, A. M. Collins, L. M. Sollid, and G. Yaari. 2019. Mosaic deletion patterns of the human antibody heavy chain gene locus shown by Bayesian haplotyping. Nature Communications 10: 628.

47. Peres, A., M. Gidoni, P. Polak, and G. Yaari. 2019. RAbHIT: R Antibody Haplotype Inference Tool. Bioinformatics 35: 4840–4842.

48. Al’Khafaji, A. M., J. T. Smith, K. V. Garimella, M. Babadi, M. Sade-Feldman, M. Gatzen, S. Sarkizova, M. A. Schwartz, V. Popic, E. M. Blaum, A. Day, M. Costello, T. Bowers, S. Gabriel, E. Banks, A. A. Philippakis, G. M. Boland, P. C. Blainey, and N. Hacohen. 2021. High-throughput RNA isoform sequencing using programmable cDNA concatenation. bioRxiv: 2021.2010.2001.462818.

49. Brochu, H. N., E. Tseng, E. Smith, M. J. Thomas, A. M. Jones, K. R. Diveley, L. Law, S. G. Hansen, L. J. Picker, M. Gale, and X. Peng. 2020. Systematic Profiling of Full-Length Ig and TCR Repertoire Diversity in Rhesus Macaque through Long Read Transcriptome Sequencing. The Journal of Immunology 204: 3434.

50. Kidd, M. J., Z. Chen, Y. Wang, K. J. Jackson, L. Zhang, S. D. Boyd, A. Z. Fire, M. M. Tanaka, B. A. Gaëta, and A. M. Collins. 2012. The Inference of Phased Haplotypes for the Immunoglobulin H Chain V Region Gene Loci by Analysis of VDJ Gene Rearrangements. The Journal of Immunology 188: 1333.

51. Kirik, U., L. Greiff, F. Levander, and M. Ohlin. 2017. Parallel antibody germline gene and haplotype analyses support the validity of immunoglobulin germline gene inference and discovery. Mol Immunol 87: 12–22.

52. Irvine, E. B., and G. Alter. 2020. Understanding the role of antibody glycosylation through the lens of severe viral and bacterial diseases. Glycobiology 30: 241–253.

53. Alter, G., T. H. M. Ottenhoff, and S. A. Joosten. 2018. Antibody glycosylation in inflammation, disease and vaccination. Seminars in Immunology 39: 102–110.

54. Collin, M. 2020. Antibody glycosylation as an immunological key in health and disease. Glycobiology 30: 200–201.

55. Plomp, R., L. R. Ruhaak, H.-W. Uh, K. R. Reiding, M. Selman, J. J. Houwing-Duistermaat, P. E. Slagboom, M. Beekman, and M. Wuhrer. 2017. Subclass-specific IgG glycosylation is associated with markers of inflammation and metabolic health. Scientific Reports 7: 12325.

56. Trampert, D. C., L. M. Hubers, S. F. J. van de Graaf, and U. Beuers. 2018. On the role of IgG4 in inflammatory conditions: lessons for IgG4-related disease. Biochimica et Biophysica Acta (BBA) - Molecular Basis of Disease 1864: 1401–1409.

