## Supplemental Tables 1-4 for "FLAIRR-seq: A novel method for single molecule resolution of near full-length immunoglobulin heavy chain repertoires"

Supplemental Table 1: FLAIRR-seq Donor Information

| Sample | Short ID | Gender | Age | Ethnicity | Smoker | Weight | Height | Viability | Blood Type |
|---|---|---|---|---|---|---|---|---|---|
| 2001430007 | 0007 | Male | 37.0 yr | Caucasian | No (Non-smoker) | 139 kg | 193 cm | 99.00 % | O negative |
| 200382201C | 2201c | Male | 32.0 yr | Caucasian | No (Non-smoker) | 87 kg | 185 cm | 94.00 % | B positive |
| 200871203C | 203c | Male | 56.0 yr | Caucasian | No (Non-smoker) | 115 kg | 165 cm | 95.00 % | A positive |
| 200381602C | 602c | Female | 24.0 yr | African American | No (Non-smoker) | 91 kg | 168 cm | 99.00 % | O positive |
| 200180705C | 705c | Male | 36.0 yr | Caucasian | No (Non-smoker) | 109 kg | 180 cm | 99.00 % | A positive |
| 2005421008 | 1008 | Female | 23.0 yr | Caucasian | No (Non-smoker) | 65 kg | 171 cm | 98.00 % | A positive |
| 2003411013 | 1013 | Male | 27.0 yr | Hispanic | No (Non-smoker) | 70 kg | 173 cm | 99.00 % | B positive |
| 2003402008 | 2008 | Female | 21.0 yr | Mixed Ethnicity | No (Non-smoker) | 57 kg | 153 cm | 99.00 % | A negative |
| 2001414002 | 4002 | Male | 27.0 yr | Caucasian | Yes (Smoker) | 104 kg | 183 cm | 92.00 % | A negative |
| 2005405001 | 5001 | Female | 56.0 yr | Asian | No (Non-smoker) | 65 kg | 160 cm | 100.00 % | B positive |
| PICR7356 | PICR7356 | Male | 41.0 yr | Non-Hispanic or Latino | N/A | N/A | N/A | N/A | N/A |
| (UofL) School of Medicine Healthy Donor | N/A | Male | 57.0 yr | Caucasian | N/A | N/A | N/A | N/A | N/A |

### Supplemental Table 2. Primers and Barcodes used for FLAIRR-seq Molecular Method

| Primer Name | Barcode 5->3 | Target 5->3 | Final Sequence 5->3 |
|---|---|---|---|
| IgG_CH3_bc1001 | CACATATCAGAGTGCG | CATGCATCACGGAGCATGAG | CACATATCAGAGTGCGCATGCATCACGGAGCATGAG |
| IgG_CH3_bc1002 | ACACACAGACTGTGAG | CATGCATCACGGAGCATGAG | ACACACAGACTGTGAGCATGCATCACGGAGCATGAG |
| IgG_CH3_bc1003 | ACACATCTCGTGAGAG | CATGCATCACGGAGCATGAG | ACACATCTCGTGAGAGCATGCATCACGGAGCATGAG |
| IgG_CH3_bc1004 | CACGCACACACGCGCG | CATGCATCACGGAGCATGAG | CACGCACACACGCGCGCATGCATCACGGAGCATGAG |
| IgG_CH3_bc1005 | CACTCGACTCTCGCGT | CATGCATCACGGAGCATGAG | CACTCGACTCTCGCGTCATGCATCACGGAGCATGAG |
| IgG_CH3_bc1006 | CATATATATCAGCTGT | CATGCATCACGGAGCATGAG | CATATATATCAGCTGTCATGCATCACGGAGCATGAG |
| IgG_CH3_bc1008 | ACAGTCGAGCGCTGCG | CATGCATCACGGAGCATGAG | ACAGTCGAGCGCTGCGCATGCATCACGGAGCATGAG |
| IgG_CH3_bc1012 | ACACTAGATCGCGTGT | CATGCATCACGGAGCATGAG | ACACTAGATCGCGTGTCATGCATCACGGAGCATGAG |
| IgM_CH4_bc1001 | CACATATCAGAGTGCG | GTCTCCCCCGTGTTCCATTC | CACATATCAGAGTGCGGTCTCCCCCGTGTTCCATTC |
| IgM_CH4_bc1002 | ACACACAGACTGTGAG | GTCTCCCCCGTGTTCCATTC | ACACACAGACTGTGAGGTCTCCCCCGTGTTCCATTC |
| IgM_CH4_bc1003 | ACACATCTCGTGAGAG | GTCTCCCCCGTGTTCCATTC | ACACATCTCGTGAGAGGTCTCCCCCGTGTTCCATTC |
| IgM_CH4_bc1004 | CACGCACACACGCGCG | GTCTCCCCCGTGTTCCATTC | CACGCACACACGCGCGGTCTCCCCCGTGTTCCATTC |
| IgM_CH4_bc1005 | CACTCGACTCTCGCGT | GTCTCCCCCGTGTTCCATTC | CACTCGACTCTCGCGTGTCTCCCCCGTGTTCCATTC |
| IgM_CH4_bc1006 | CATATATATCAGCTGT | GTCTCCCCCGTGTTCCATTC | CATATATATCAGCTGTGTCTCCCCCGTGTTCCATTC |
| IgM_CH4_bc1008 | ACAGTCGAGCGCTGCG | GTCTCCCCCGTGTTCCATTC | ACAGTCGAGCGCTGCGGTCTCCCCCGTGTTCCATTC |
| IgM_CH4_bc1012 | ACACTAGATCGCGTGT | GTCTCCCCCGTGTTCCATTC | ACACTAGATCGCGTGTGTCTCCCCCGTGTTCCATTC |
| TSO_UMI | 5’- AAGCAGUGGTAUCAACGCAGAGUNNNNUNNNNUNNNNUCTTrGrGrG -3’ | | |

Supplemental Table 3: AIRR-seq pRESTO and Change-O Pipeline Read Counts

| Sample | Start R1 | Start R2 | Quality Filter R1 | Quality Filter R2 | Length Filter R1 | Length Filter R2 | Mask Primer Pair R1 IGM | Mask Primer Pair R1 IGG | Mask Primer Pair R2 | Consensus R1 IGM | Consensus R1 IGG |
|---|---|---|---|---|---|---|---|---|---|---|---|
| 0007 | 841627 | 841627 | 840447 | 824203 | 809108 | 754640 | 424261 | 320692 | 744953 | 42691 | 19294 |
| 201c | 1509063 | 1509063 | 1508821 | 1486250 | 1443879 | 1359277 | 818708 | 527372 | 1346080 | 98872 | 33489 |
| 203c | 1006965 | 1006965 | 1005528 | 987965 | 966698 | 900107 | 593540 | 298804 | 892344 | 80256 | 31183 |
| 602c | 1275902 | 1275902 | 1274218 | 1253423 | 1229554 | 1149372 | 512249 | 627449 | 1139698 | 40180 | 54527 |
| 705c | 1113660 | 1113660 | 1113251 | 1094451 | 1065379 | 994472 | 441845 | 542531 | 984376 | 62271 | 21622 |
| 1008 | 1310350 | 1310350 | 1308354 | 1279670 | 1257122 | 1147973 | 563435 | 575400 | 1138835 | 85370 | 93698 |
| 1013 | 1158222 | 1158222 | 1157870 | 1137860 | 1115429 | 1045281 | 510527 | 524412 | 1034939 | 40840 | 32531 |
| 2008 | 835928 | 835928 | 835584 | 818130 | 804971 | 742595 | 343383 | 392596 | 735979 | 40470 | 27887 |
| 4002 | 632272 | 632272 | 632128 | 620509 | 603814 | 546881 | 326911 | 172689 | 499600 | 55138 | 12033 |
| 5001 | 695884 | 695884 | 695568 | 685207 | 657394 | 605206 | 323743 | 237184 | 560927 | 37088 | 18169 |

| Sample | Consensus Pair R1 IGM | Consensus Pair R1 IGG | Consensus Pair R2 | Assemble IGM | Assemble IGG | Collapse IGM | Collapse IGG | Split IGM | Split IGG | IG Blast IGM | IG Blast IGG |
|---|---|---|---|---|---|---|---|---|---|---|---|
| 0007 | 42691 | 19294 | 54036 | 30831 | 15217 | 12777 | 6281 | 9363 | 3205 | 9154 | 3164 |
| 201c | 98872 | 33489 | 105587 | 66722 | 25015 | 26850 | 10031 | 18353 | 4639 | 17888 | 4568 |
| 203c | 80256 | 31183 | 88067 | 54483 | 23085 | 24623 | 9881 | 18712 | 5864 | 18189 | 5793 |
| 602c | 40180 | 54527 | 79316 | 29949 | 38078 | 12306 | 16553 | 8584 | 7851 | 8378 | 7736 |
| 705c | 62271 | 21622 | 68358 | 42597 | 15358 | 19484 | 4621 | 14700 | 1491 | 14273 | 1362 |
| 1008 | 85370 | 93698 | 126939 | 55556 | 56848 | 20264 | 26612 | 13022 | 12449 | 12710 | 12217 |
| 1013 | 40840 | 32531 | 65769 | 30765 | 25250 | 12769 | 10135 | 9201 | 5192 | 9012 | 5027 |
| 2008 | 40470 | 27887 | 58854 | 30069 | 21681 | 11608 | 8762 | 8734 | 3965 | 8551 | 3900 |
| 4002 | 55138 | 12033 | 59929 | 42915 | 8949 | 18411 | 3549 | 11845 | 1145 | 11607 | 1112 |
| 5001 | 37088 | 18169 | 51206 | 27927 | 13930 | 10018 | 4991 | 6464 | 1937 | 6314 | 1885 |

Supplemental Table 4: FLAIRR-seq pRESTO and Changeo-O Pipeline Read Counts

| Sample | Quality Filter | Mask Primer | Mask Primer IGG | Consensus IGG | Collapse IGG | Split IGG | IGG Blast |
|---|---|---|---|---|---|---|---|
| 0007 | 181109 | 169240 | 169240 | 17881 | 12794 | 6641 | 4880 |
| 201c | 221301 | 210050 | 210050 | 36053 | 26043 | 13057 | 10819 |
| 203c | 475703 | 467231 | 467231 | 14923 | 10261 | 4546 | 3778 |
| 602c | 344906 | 332644 | 332644 | 33903 | 23979 | 10750 | 9225 |
| 705c | 279601 | 272045 | 272045 | 18627 | 12913 | 5038 | 2328 |
| 1008 | 238820 | 232762 | 232762 | 57020 | 41061 | 20314 | 18235 |
| 1013 | 396322 | 388655 | 388655 | 27871 | 20603 | 10891 | 7487 |
| 2008 | 418906 | 407961 | 407961 | 16441 | 11081 | 5706 | 4258 |
| 4002 | 398838 | 374062 | 374062 | 19692 | 15091 | 6830 | 5377 |
| 5001 | 313843 | 306159 | 306159 | 42845 | 25359 | 14970 | 12146 |
| **Sample** | **Quality Filter** | **Mask Primer** | **Mask Primer IGM** | **Consensus IGM** | **Collapse IGM** | **Split IGM** | **IGM Blast** |
| 0007 | 331380 | 307398 | 307398 | 46384 | 40581 | 18178 | 14926 |
| 201c | 148355 | 140380 | 140380 | 98069 | 86061 | 17095 | 12803 |
| 203c | 509536 | 506280 | 506280 | 37527 | 31679 | 12476 | 10636 |
| 602c | 273103 | 268267 | 268267 | 48897 | 43704 | 19601 | 16406 |
| 705c | 435391 | 427305 | 427305 | 79251 | 74897 | 21045 | 17417 |
| 1008 | 181063 | 171100 | 171100 | 47531 | 40075 | 18353 | 15284 |
| 1013 | 262617 | 258749 | 258749 | 52508 | 46827 | 21303 | 17812 |
| 2008 | 502354 | 498342 | 498342 | 47596 | 39149 | 16393 | 14132 |
| 4002 | 489287 | 476829 | 476829 | 91732 | 87654 | 26195 | 22379 |
| 5001 | 171688 | 163545 | 163545 | 18944 | 16004 | 8102 | 6164 |
